# Dynamic X-chromosomal reactivation enhances female brain resilience

**DOI:** 10.1101/2023.06.17.545424

**Authors:** S Käseberg, M Bertin, R Menon, E Gabassi, H Todorov, S Frank, H Brennenstuhl, B Lohrer, J Winter, J Krummeich, J Winkler, B Winner, E Weis, D Hartwich, S Diederich, K Luck, S Gerber, P Lunt, B Berninger, S Falk, S Schweiger, M Karow

## Abstract

While random X-chromosome inactivation in female cells of placental mammalians silences one allele of the majority of X-chromosomal genes, a considerable fraction is only incompletely and variably inactivated resulting in a tissue-specific pattern of mono- and biallelic expression. Here we used clonal human female induced pluripotent stem cells (iPSCs) allowing to trace the (in)activation status of the two X-chromosomes individually along neural differentiation trajectories. We discovered X-chromosome-wide locus- and lineage-specific dynamic usage of the two X-chromosomal alleles in female cells induced by differentiation. Leveraging iPSCs derived from patients with an X-linked neurodevelopmental disorder, we demonstrate that activation of alleles on the inactive X-chromosome can exert protective effects on the manifestation of disease phenotypes in female neural cells and tissue. Taken together, our data demonstrate that alleles on the inactive X-chromosome can serve as a critical reservoir reactivated during differentiation, thereby enhancing resilience of female neural tissue.

## Introduction

In somatic cells of female eutherian mammals one of the two X-chromosomes is randomly inactivated leading to a silent gene reservoir (Sahakyan et al., 2018). Early during embryonic development X-chromosomal genes undergo inactivation at different timepoints during tissue differentiation (Patrat et al., 2009; Petropoulos et al., 2016). Some of the genes on the inactive X-chromosome, however, have been found to escape the inactivation process (Berletch et al., 2011; Carrel and Willard, 2005; Fang et al., 2019). In addition to constitutive escapees, which escape X-chromosome inactivation (XCI) in all cells and tissues, a similar number of genes escape inactivation facultatively. Facultative entails that biallelic expression is observed only in selected tissues and is heterogenous between individuals (Berletch et al., 2015; Tukiainen et al., 2017; Zhang et al., 2013). Both, XCI as well as escapism show considerable differences between eutherian mammalian species (Genolet et al., 2021; Okamoto et al., 2021; Patrat et al., 2020; Petropoulos *et al*., 2016), pointing towards different regulatory mechanisms and highlighting the importance of species-specific model systems. Incomplete XCI results in biallelic expression and in humans frequently in a female expression bias, i.e. higher expression levels of the same gene in female tissue compared to the corresponding male tissue (Oliva et al., 2020; Tukiainen *et al*., 2017). However, there is hardly any experimental data shedding light on the developmental timeline of X-inactivation and escapism and the question how facultative escape emerges in human cells. Taking advantage of the clonal nature and the retention of the X-chromosome activation status in human induced pluripotent stem cells (iPSCs) (Tchieu et al., 2010), we here monitored XCI and escapism throughout tissue differentiation from a defined point-of-origin. We observed dynamic reactivation and late-silencing of alleles on the inactive X-chromosomes induced by neural differentiation. Strikingly, many of these reactivating genes are linked to neurodevelopmental disorders (NDDs).

NDDs exhibit an intriguing sex bias with females being less often and less severely affected than males (Pearse and Young-Pearse, 2019). While NDD associated genes are evenly distributed across all autosomes, they are overrepresented on the X-chromosome (Leblond et al., 2021) suggesting a particularly important role of the X-chromosome in the development of NDDs. In males, most X-linked genes are hemizygous, i.e., males only carry the allele inherited from the mother. In contrast, random XCI in female cells results in a mosaic expression of X-linked genes in female tissue, substantially influencing phenotype development of X-linked gene defects. However, hemizygosity of X-linked genes in males and random XCI in females only partially explain the fact that females are disproportionally less affected by NDDs (Brand et al., 2021; Jacquemont et al., 2014; Ross et al., 2005). We hypothesized that dynamic X-chromosome reactivation is part of a protective shield in female brains reducing frequency and severity of NDDs in girls. To test this hypothesis, we employed an iPSC-based brain organoid model of Opitz BBB/G syndrome (OS), an X-linked NDD caused by mutations in the *MID1* gene. While brain organoids derived from a male OS patient exhibited a dramatic increase in neural progenitors at the expense of differentiated neurons, we found that female organoids with the same mutation on their active X-chromosome showed a markedly milder, intermediate phenotype. To interrogate the influence of the allele on the inactive X-chromosome, we employed CRISPR/Cas9-mediated introduction of the same loss-of-function allele on both the active and inactive X-chromosome. Strikingly, brain organoids derived from this homozygous female OS iPSC line exhibited a phenotype similar to the hemizygous male organoids. This underscores that the allele on the inactive X-chromosome substantially influences phenotype development.

Taken together, we here dissected the temporal usage of the two X-chromosomes along neural differentiation trajectories and thereby uncloak reactivation of alleles on the inactive X-chromosome as a mechanistic basis resulting in facultative escapism. Furthermore, we show that this fine-tuned symphony of dynamic biallelic X-chromosomal gene expression has substantial protective effects contributing to female brain resilience.

## Results

### X-reactivation during differentiation

To trace the X-activation status along differentiation trajectories, we took advantage of the clonal nature of human iPSCs maintaining the X-inactivation status of the cell of origin (Tchieu *et al*., 2010) and leveraged heterozygous sequence variants on the X-chromosome. This enabled us to follow the usage of individual alleles on the X-chromosome along the temporal axis from a defined point-of-origin (iPSCs) during differentiation and allowed to address the “When and Where” of X-inactivation and escapism. Female fibroblasts were reprogrammed into iPSCs and sequence variants that distinguished the two X-chromosomes were used for clonal selection (Figures S1A-D). Using bulk RNA-sequencing of iPSCs, neural stem and progenitor cells (NPCs), and neurons, followed by allele-specific expression analyses we performed a global inspection of the allele usage of X-linked genes (Figure 1A). Cluster and principal component analysis of the bulk RNA-sequencing data proved a clear distinction between iPSCs, NPCs, and neurons based on their allelic ratio profiles (Figures S1E).

**Figure 1.**
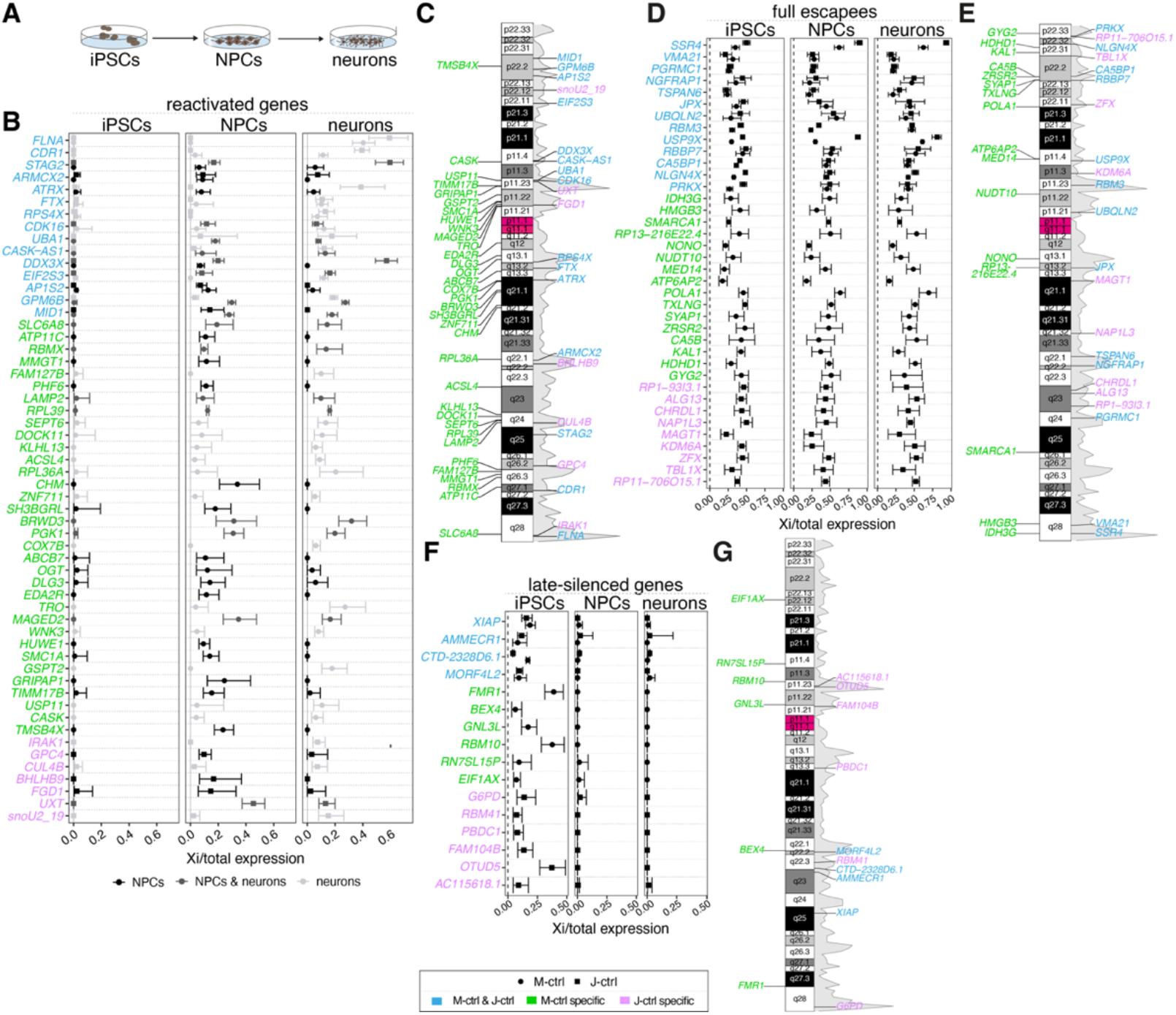
Reactivation of X-linked genes during neuronal differentiation in female cells. **A,** Experimental scheme depicting neural 2D differentiation. **B,** Scatter plots showing the ratio of expression from the inactive X- chromosome per total allelic mRNA expression for genes biallelically expressed in NPCs and/or neurons but not in iPSCs. **C,** Ideogram of the X-chromosome showing the cytogenetic location of reactivated genes in the M-ctrl, J-ctrl or both cell lines. The gray area represents the gene density along the X-chromosome using 1 Mbp genomic windows. **D**, Scatter plots showing the ratio of expression from the inactive X-chromosome per total allelic mRNA expression for escape genes, i.e., genes with biallelic expression in iPSCs and throughout the differentiation paradigm. **E**, Ideogram of the X-chromosome showing the cytogenetic location of full escapees in the M-ctrl, J- ctrl or both cell lines. The gray area represents the gene density along the X-chromosome using 1 Mbp genomic windows. **F**, Scatter plots showing the ratio of expression from the inactive X-chromosome per total allelic mRNA expression for late-silenced genes, i.e., genes with biallelic expression in iPSCs but not in NPCs or neurons. **G**, Ideogram of the X-chromosome showing the cytogenetic location of late-silenced genes in the M-ctrl, J-ctrl or both cell lines. The gray area represents the gene density along the X-chromosome using 1 Mbp genomic windows. **B-G**, M-ctrl iPSCs: *n=*6, J-ctrl iPSCs: *n=*8, NPCs & neurons of M-ctrl and J-ctrl: n=4. Error bars indicate the 99% confidence intervals.

A total of 49 (M-ctrl) and 22 (J-ctrl) genes were monoallelically expressed in iPSCs but exhibited differentiation-dependent reactivation (reactivating genes) resulting in biallelic expression in either NPCs, neurons, or both cell types (Figures 1B, C). In addition to gene reactivation, we detected 29 (M-ctrl) and 22 (J-ctrl) genes that were expressed biallelically in iPSCs, NPCs, and neurons (full escapees, Figure 1D, E) and 10 (M-ctrl) and 10 (J-ctrl) genes that switched from biallelic expression in iPSCs to monoallelic expression in NPCs and neurons (late-silencing genes, Figure 1F, G). While full escapees, as expected, showed similar expression levels from both alleles (biallelic expression level around 0.5), reactivated and late-silenced genes exhibited lower levels of average expression from the inactive allele in the respective cell type (Figure 2A). Importantly, all three gene categories showed no correlation between expression level and biallelic expression (Figure S1F). Quantification of *XIST* and *XACT* expression and the biallelic expression ratio between the X-chromosomes and autosomes (X:A allelic ratio), demonstrated proper XCI without erosion (Bar et al., 2019) in the iPSC clones and their derivatives used here (Figures S1G, H). For validation we used Quantification of Allele-Specific Expression by Pyrosequencing (QUASEP) assays to quantify allelic expression of selected genes based on expressed single nucleotide polymorphisms (SNPs). While *CA5B*, a gene known to constitutively escape XCI (Garieri et al., 2018), was expressed biallelically in all cells tested, including iPSCs, *ZNF185* showed purely monoallelic expression throughout neuronal differentiation. In contrast *MID1* as well as *GPM6B*, two genes located on the short arm of the X-chromosome in close vicinity (Figure 1C), were reactivated in NPCs and neurons indicating locus-specific reactivation (Figures S1I-N). To further confirm reactivation of alleles on the inactive X-chromosome we leveraged the traceability of a 4-base pair (bp) deletion in the *MID1* gene on the inactive X-chromosome (M-ctrl) and derived and engineered iPSC lines with the same sequence variant on the active X-chromosome (M-OS/het, J-OS/het; Figure S1A-D, O). Using allele-specific RT-PCR, substantial reactivation of the *MID1* allele on the inactive X-chromosome was found in both, NPCs and neurons derived from the M-ctrl, M-OS/het and J-OS/het lines while the respective iPSCs showed strictly monoallelic expression (Figure S1P). Western blot analysis revealed that the reactivation of the wildtype allele from the inactive X-chromosome resulted in the expression of MID1 wildtype protein in the M-OS/het and J-OS/het NPCs, while no MID1 protein expression was detected in the corresponding iPSCs (Figures S1Q, R). These data uncover that in addition to purely monoallelic and full escapees, additional categories of X-chromosomal genes exist that can be reactivated or silenced during neuronal differentiation thereby provoking facultative escapism.

**Figure 2.**
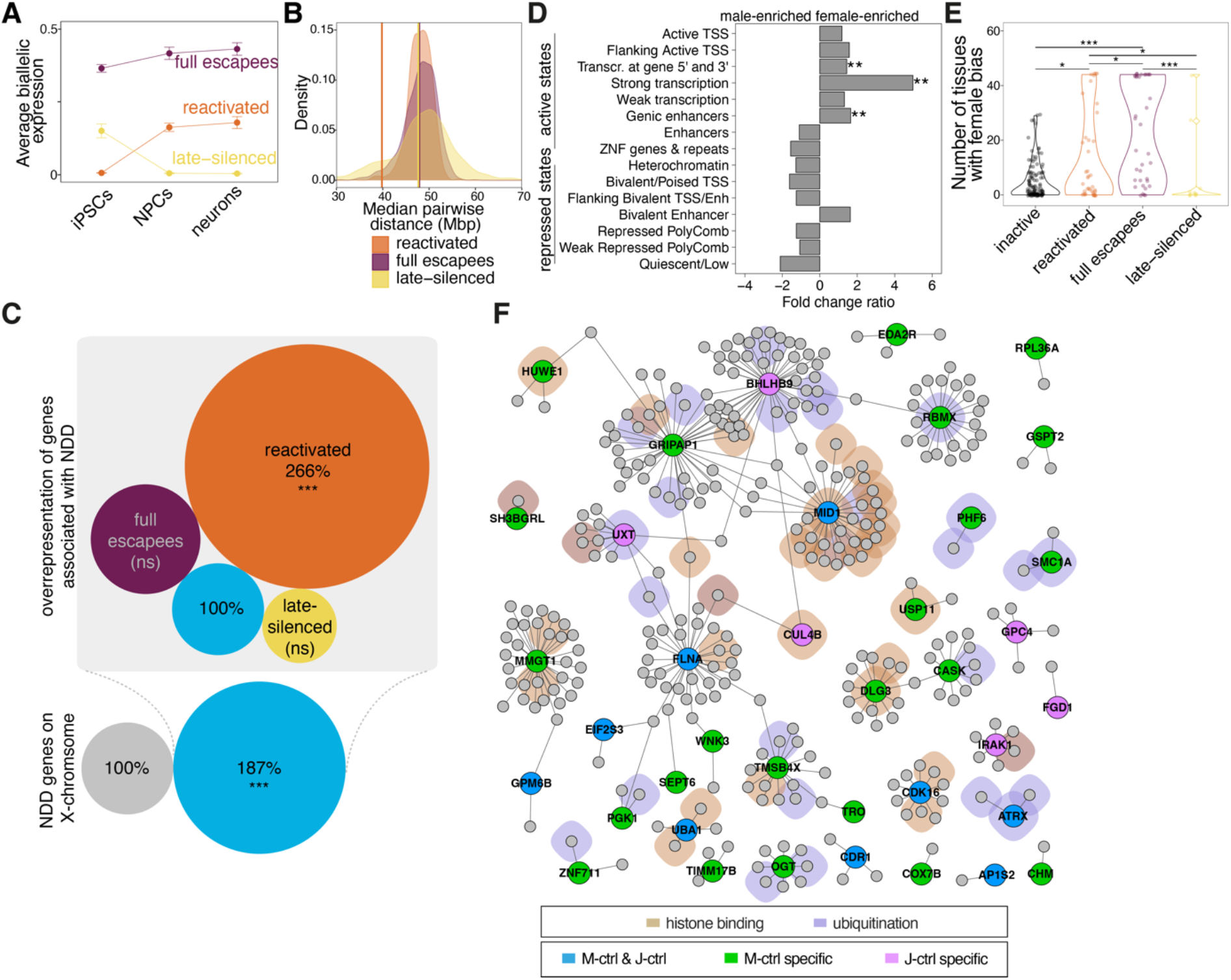
Genomic organization and characterization of biallelically expressed genes. **A,** Average biallelic expression levels (proportion of expression from Xi) for the three types of biallelically expressed genes in iPSCs, NPCs and neurons. Biallelic expression levels were averaged across all genes in the respective category and across the M-ctrl and J-ctrl lines. Error bars indicate standard error. **B**, Density plot showing the background distribution of 1000 sets of randomly selected X-linked genes. The different colors indicate varying sample sizes in the groups of reactivated, escape and late-silenced genes. Vertical lines show the median pairwise distance between reactivated, escape or late-silenced genes respectively. Note that the reactivated but not full escape or late-silenced genes localize closer to each other than expected by chance. *P*(reactivated)=7.0*10^-4^, *P*(full escapee)=4.8*10^-1^, *P*(late-silenced)=4.6*10^-1^. *P*-values were calculated using the cumulative distribution function of the normal distribution. **C,** Plot showing the overrepresentation of the NDD genes on the X-chromosome (187% more than expected; *P*<10^-6^). While in full escapees and late-silenced genes no further enrichment of X-linked NDD genes was detected (escapee: 119%, *P*=0.37; late-silenced: 81%; *P*=0.73), in reactivated genes an intriguing accumulation beyond the overrepresentation of NDD genes on the X-chromosome was observed (266% more than expected; *P*=10^-6^). *P-*values were determined by the cumulative distribution function. **D,** Bar plot showing the fold change ratio of chromatin state enrichment (Roadmap Epigenomics *et al*., 2015) between reactivated and inactive genes in female relative to male fetal brain tissue. Positive values indicate higher chances for the corresponding state to be enriched in reactivated genes in females. Exact *P*-values (top to bottom): 6.8*10^-3^, 4.7*10^-3^, 6.8*10^-3^. **E,** Violin plot showing the distribution of the number of tissues showing a female-biased gene expression for the different categories of biallelically expressed as well as inactive genes, exact *P*-values (top to bottom and left to right): 6.7*10^-7^, 3.4*10^-2^, 1.7*10^-2^, 3.4*10^-2^, 1.5*10^-4^. **F,** A protein-protein interaction (PPI) network of protein-coding reactivated genes based on genes expressed in NPCs and neurons. Interactions between the PPI-partners not involving reactivated genes are not shown. GO analysis revealed a significant overrepresentation of processes related to chromatin/histone binding and protein ubiquitination processes. Reactivated genes and their interaction partners that are involved in these processes are indicated on the network.

In contrast to full escapees and late-silencing genes, reactivating genes were not randomly distributed but rather clustered together on the X-chromosome as indicated by significantly lower pairwise distances than expected by chance (Figure 2B) providing evidence for a spatial organization of reactivated genes along the X-chromosome. Strikingly, we observed a reactivation cluster on Xp11.2 (Figure 1C) which is a region heavily involved in X-linked intellectual disability (Moey et al., 2016; Qiao et al., 2008). This raised the question whether X-reactivation is a more general component associated with NDDs. Indeed, amongst the reactivating genes, but not full escapees and late-silencing genes, we discovered an enrichment of NDD associated genes beyond the expected overrepresentation on the X-chromosome (Brand *et al*., 2021) (Figure 2C). Our data suggests that in particular genes associated with neurodevelopment and NDDs were evolutionary endowed with dynamic X-reactivation as a regulatory mechanism.

By employing the *15-state chromatin model (Roadmap Epigenomics et al., 2015)*, we identified a distinct epigenetic pattern for reactivating genes that was more associated with active transcription and genic enhancer states, while genes for which we did not detect biallelic expression showed a tendency for enrichment of repressed chromatin states (Figure S1S). Importantly, the chances for the active states (‘Transcription at gene 5’and 3’’, ‘Strong transcription’, ‘Genic enhancers’) being enriched in reactivating versus inactive genes is significantly higher in fetal female brain tissue compared to male (Figure 2D). These epigenetic differences between reactivating and non-reactivating genes indicate that changes on the chromatin level allow reactivation. Biallelic expression of X-chromosomal genes in female cells could lead to female expression bias. By analyzing expression pattern in 44 different tissues (Oliva *et al*., 2020), we found a gradual increase in the number of tissues showing a female expression bias from late-silencing and solely monoallelic towards reactivating and full escapees (Figure 2E), suggesting that the genes here detected as biallelically expressed may also undergo dynamic X-reactivation along non neural lineages. To investigate the molecular framework the reactivating genes are contributing to, we constructed a protein-protein interaction network based on genes expressed in NPCs and neurons (Figure 2F). A gene ontology (GO) enrichment analysis revealed an overrepresentation of terms associated with epigenetic regulatory processes such as histone binding and, in parts, protein ubiquitination (Figure S1T, Table S1) highlighting the potential global impact on gene and protein function by the reactivation of X-linked genes.

### Dynamic expression from the inactive X-chromosome

To obtain a fine-grained picture of X-chromosomal allele-usage across developmental trajectories, we generated brain organoids from the above-mentioned iPSC lines (Figure S1A, O) and performed scRNA-seq (Figure S2A-E). Force directed graph embedding and annotation of the distinct lineages within the dataset utilizing lineage-specific marker genes, revealed an eminent hindbrain regional identity of the neural tissue, as well as non-neural mesenchymal clusters (Figures 3A, S2F). All samples contributed to a similar extent to all lineages and clusters (Figure S2G). We first determined heterozygously expressed and sequenced X-chromosomal SNPs in the scRNA-seq dataset, constructed a variant-aware reference genome and remapped the raw sequencing data to it. We detected a total of 26 X-linked genes with a traceable SNP, i.e. a SNP located within 150 bp upstream of the polyA-tail. Albeit the different cellular sources of the samples and the employed techniques, several of these genes overlapped with the genes detected in the bulk RNA-seq data (Figures 1, S2H). While the bulk RNA-seq approach was restricted to iPSCs, NPCs and neurons, this approach allowed us to follow the extent of biallelic expression of individual genes on a single cell level over multiple differentiation lineages.

**Figure 3.**
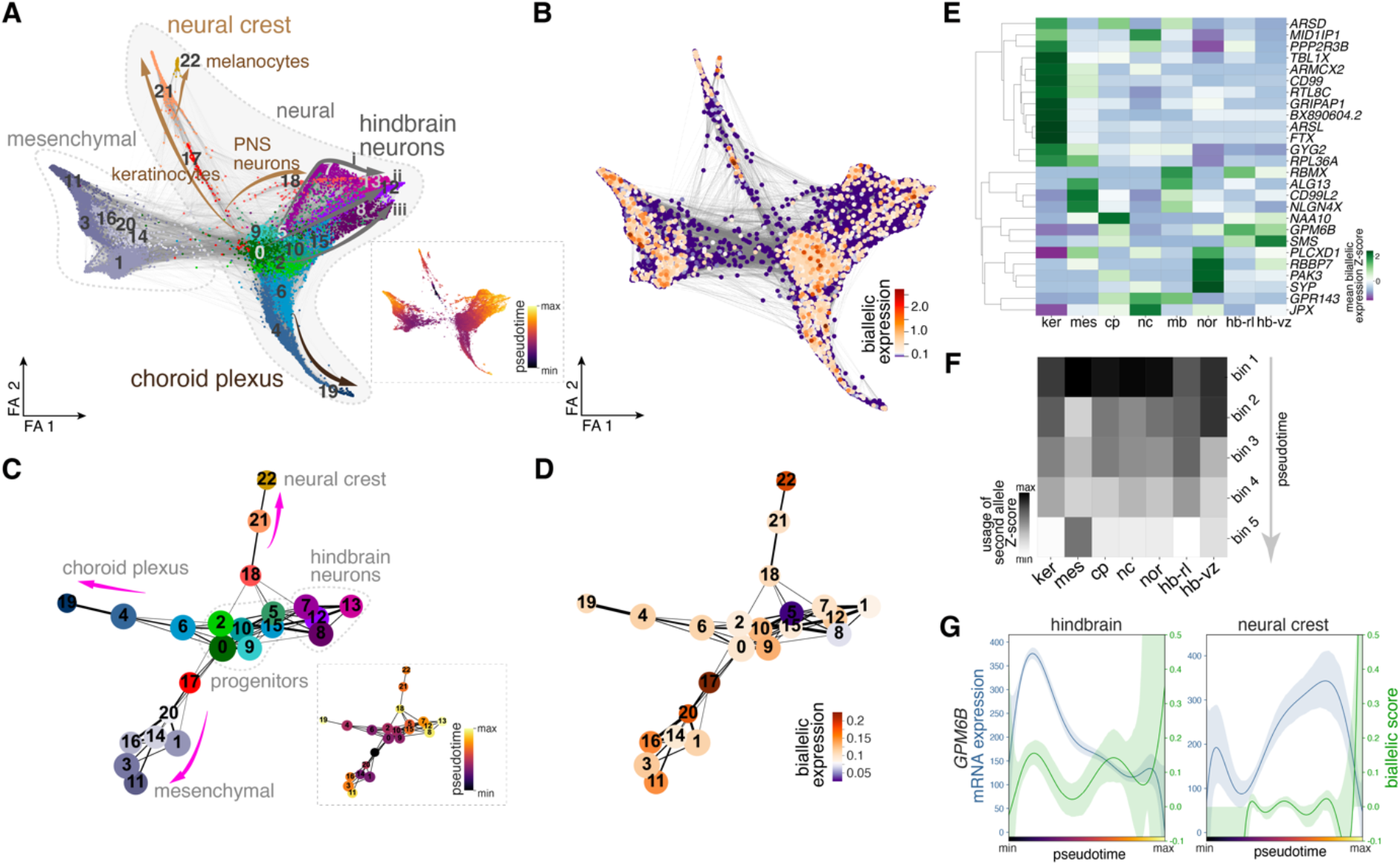
Dynamic biallelic expression of X-chromosomal genes across differentiation trajectories. **A,** Force directed graph embedding showing clusters and annotated lineages as further shown in PAGA graph abstraction in **C**. RNA-velocity based pseudotime is indicated in the dashed box. Number of cells total: 16331, M-ctrl: 2236; M-OS/het: 6326; J-ctrl: 2113; J-OS/het: 1301; J-OS/hom: 4355. **B,** Degree of biallelic expression of X-linked genes projected onto the force directed graph. **A, B**, FA refers to force atlas. **C,** PAGA plot showing lineage trajectories connecting clusters and the corresponding RNA-velocity derived pseudotime (dashed box). **D,** Cluster based PAGA plot showing the biallelic expression of X-chromosomal genes reveals varying overall biallelic expression between clusters (indicated with numbers) and along trajectories. **E,** Heatmap depicting the Z-score of biallelic expression of the 26 genes detected to be biallelically expressed in distinct differentiation lineages. **F,** Heatmap showing the average biallelic expression of X-chromosomal genes along distinct lineages binned in 5 equally sized pseudotemporal bins along the differentiation pseudotime. **E, F,** ker (keratinocytes), mes (mesenchymal), nc (neural crest), mb (midbrain), nor (noradrenergic), hb-rl (hindbrain-rhombic lip), hb-vz (hindbrain-ventricular zone). **G,** Pseudotemporally resolved lineage-specific expression dynamics of *GPM6B* mRNA and biallelic expression score in hindbrain (left panel) and neural crest (right panel) along their respective differentiation trajectories.

Visualizing global biallelic expression on a single cell level (Figure 3B) and on a cluster level (Figures 3C, D) using the PAGA graph abstraction (Wolf et al., 2019) showed that all lineages exhibited biallelic expression but that the degree differed between lineages and along differentiation trajectories. Individual genes exhibited strong differences in the extent of biallelic expression with some genes showing broad usage of the second allele across all lineages (e.g. *RPL36A*) while for others, biallelic expression was more restricted (e.g. *GPM6B*) and for some even limited to single lineages (e.g. *PAK3*) (Figure S2H). We next grouped the cells according to their lineage and calculated the level of mean biallelic expression of the individual genes. The resulting heatmap revealed a lineage-specific biallelic expression pattern supporting the notion of an actively controlled process (Figure 3E). To temporally resolve X-chromosome usage within individual lineages we used RNA velocity-based pseudotime estimation (Figures 3A, C) (Bergen et al., 2020; La Manno et al., 2018) and binned each lineage in 5 equally sized pseudotemporal bins. For each lineage we calculated the normalized usage of the alleles on the inactive X-chromosome. This approach unambiguously revealed a high temporal dynamic in the usage of the alleles on the inactive X-chromosome along differentiation and a substantial variability between individual differentiation trajectories (Figure 3F). To rule out that the degree of biallelic expression solely depends on the rate of transcription we generated a dot-plot showing for each of the 26 biallelically expressed genes in every cluster the average mRNA expression versus the mean biallelic expression. No correlation between the degree of biallelic expression and the expression level per se was detected (Figure S3A). This notion was further supported by plotting for each gene the amount of mRNA expression and the degree of biallelic expression over the course of differentiation (Figure S3B). In both approaches no general dependency between usage of the allele on the inactive X-chromosome and the transcription level could be discerned. Plotting transcription and biallelic expression of *GPM6B*, a gene that was also identified as reactivating in the bulk RNA-seq data, along distinct differentiation trajectories (hindbrain and neural crest) revealed that the same gene with similar expression levels in both lineages can show different biallelic expression levels and dynamics (Figure 3G). In summary, these data show a temporally dynamic and lineage-specific usage of the alleles on the inactive X-chromosome during differentiation.

### Impact of X-reactivation on phenotype dimorphism

The accumulation of NDD associated genes in the group of reactivating genes suggests an influence of X-chromosomal reactivation on the pathogenesis of NDDs. In order to analyze if X-chromosomal reactivation has an influence on the phenotypic presentation of X-linked NDDs we focused our attention on Opitz BBB/G syndrome (OS) (Quaderi et al., 1997; So et al., 2005; Winter et al., 2016). OS is an NDD caused by mutations in the *MID1* gene, a gene detected as reactivating in both cell lines, M-ctrl and J-ctrl (Figures 1B, C). Affected patients present with a variable set of neurodevelopmental aberrations including hypoplasia of the cerebellar vermis (Schweiger et al., 1999). Moreover, females present with milder phenotypes than males (So *et al*., 2005). To test if reactivation of *MID1* in female cells impacts on the development of phenotypic dimorphism of OS, we compared brain organoids derived from female cells (Figures S1A, O) with organoids derived from male cells. The male iPSCs were established from a male fetus affected with OS (Figure S4B) (M-OS/male) hemizygously carrying the same 4-bp deletion in *MID1* (Figure S4A) that was present in the female OS lines. Both, M-ctrl and M-OS/het were derived from the affected heterozygous mother of the male fetus. While in the M-OS/het iPSC line the 4-bp deletion was present on the active X-chromosome, in the M-ctrl iPSC line the 4-bp deletion was silenced on the inactive X-chromosome and the wildtype allele of *MID1* was expressed (Figures S1A, C, O). In addition, we engineered a set of female iPSC lines from an unrelated control female (J) carrying wildtype *MID1* (J-ctrl) on both chromosomes. Using CRISPR/Cas9 we introduced the 4-bp deletion either heterozygously on the active X-chromosome (J-OS/het) or homozygously on both *MID1* alleles (J-OS/hom) (Figure S1A, O). While qRT-PCR in the OS iPSCs proved stability of the mutant mRNA (Figure S4C), MID1 protein could not be detected in the mutant OS iPSC lines (Figure S4D) suggesting either instability or aggregation of the mutant MID1 protein (Schweiger *et al*., 1999).

To assess how the 4-bp loss-of-function mutation of *MID1* affects early human brain development, we analyzed brain organoids with a hindbrain identity from ctrl and OS iPSC lines. Inspection of the cellular organization of the brain organoids at day 30 (d30) revealed an increase in the area covered by NPC-containing ventricular zone-like structures (VZLS) (Figures 4A, S4E) while the cell density remained unchanged (Figure S4F). Quantification of the cellular composition of brain organoids (d30 and d50) showed an increase of SOX2^+^ NPCs and a concomitant decrease in MAP2^+^ neurons within *MID1* mutant organoids (M-OS/het, M-OS/male, J-OS/het, J-OS/hom) compared to *MID1* wildtype organoids (M-ctrl, J-ctrl, Figures 4B-D, S4G-J). CRISPR/Cas9-mediated repair of the *MID1* mutation in the male cells (M-OS/maleR) led to a rescue of the phenotype (Figures 4A-C, S4E, G) proving MID1 dependency of the phenotypic effects.

**Figure 4.**
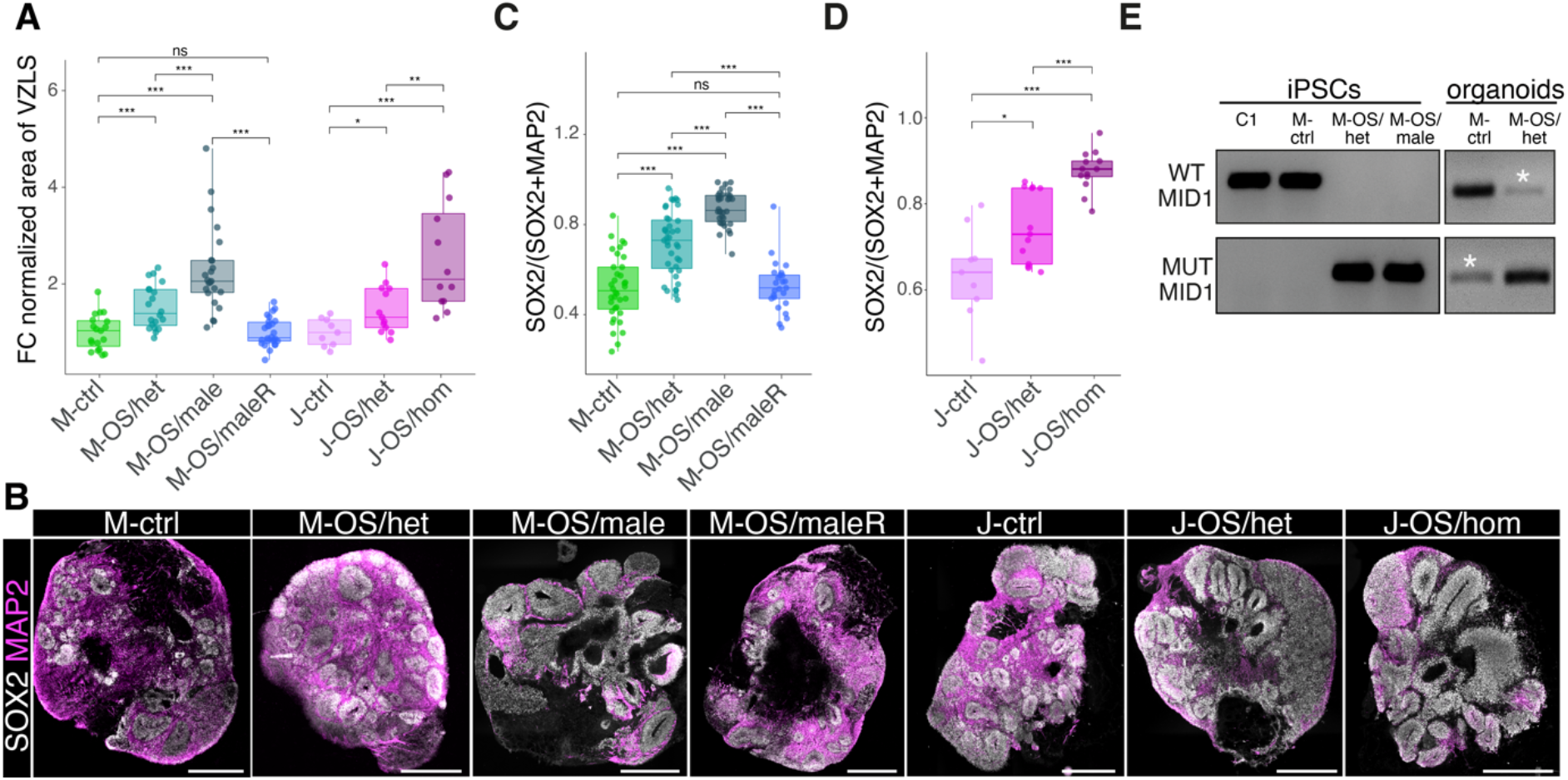
Intermediate phenotype in female *MID1* mutant brain organoids. **A,** Box- and jitter plot depicting the fold change of brain organoid area organized as VLZS compared to the respective ctrl line (M-ctrl, *n=*19; M- OS/het, *n=*20; M-OS/male, *n=*23; M-OS/maleR, *n=*26; J-ctrl, *n=*9; J-OS/het, *n=*13; J-OS/hom, *n=*12; exact *P* values (top to bottom): 0.83, 9.7*10^-4^, 4.5*10^-3^, 2.0*10^-8^, 2.7*10^-5^, 7.0*10^-4^, 2.5*10^-2^, 2.0*10^-10^). **B,** Immunostaining of organoid sections showing SOX2 (white) and MAP2 (magenta). Scale bars, 100 µm. **C, D** Quantification of the fraction of SOX2+ area per total neural area (SOX2+ or MAP2+ area) in d30 organoids. **C,** (M-ctrl, *n=*36; M-OS/het, *n=*39; M-OS/male, *n=*34; M-OS/maleR, *n=*25; exact *P* values (top to bottom): 7.5*10^-7^, 0.71, 1.4*10^-13^, 2.8*10^-7^, <2.2*10^-16^, 1.3*10^-7^). **D,** (J-ctrl, *n=*9; J-OS/het, *n=*13; J-OS/hom, *n=*13; exact *P* values (top to bottom): 3.7*10^-5^, 8.0*10^-6^, 2.1*10^-2^). **A, C, D,** boxplots show median, quartiles (box), and range (whiskers), dots represent individual organoids; two-sided Wilcoxon rank sum test; *\*P*<0.05, *\*\*P*<0.01, *\*\*\*P*<0.001, ns *P*>0.05. **E,** Allele-specific RT-PCR revealing reactivation (*) of the inactive *MID1* allele in brain organoids.

Strikingly, the brain organoids derived from the heterozygous female iPSC lines, M-OS/het and J-OS/het, exhibited a less severe phenotype compared to M-OS/male (Figure 4A, B, D). While this is reminiscent to the clinical manifestations, in our paradigm using clonal iPSC lines with a defined and stable X-(in)activation status random X-inactivation cannot contribute to the amelioration of the phenotype in female tissue. In agreement with the bulk RNA-seq data, allele specific RT-PCR showed reactivation of the *MID1* gene from the inactive X-chromosome in brain organoids (Figure 4E). To verify the influence of the wildtype *MID1* allele on the inactive X-chromosome in heterozygous female OS cells we took advantage of the female iPSCs homozygous for the 4-bp deletion (J-OS/hom). Brain organoids derived from this line showed a phenotype comparable to the male OS organoids (Figures 4A, B, D, S4E, G). These data underscore that the presence of a wildtype allele on the inactive X-chromosome in heterozygous mutant female cells curtails the severity of the disease phenotype through reactivation of the allele on the inactive X-chromosome.

### Dissecting the cellular and molecular underpinnings of the OS phenotype

To identify the molecular signatures underlying the gradual OS phenotypes, we focused our attention on the neural lineages. Within the neural tissue we found cells with molecular signatures of neural crest, choroid plexus, midbrain, and hindbrain lineages (Figures 5A, S5A). This hindbrain organoid model recapitulated the major lineages of the developing cerebellum, formation of which is disturbed in OS (MacDonald et al., 1993). NPC-to-neuron trajectories producing three distinct neurotransmitter identities were discriminated: rhombic lip progenitors (*ATOH1, LHX9*) giving rise to glutamatergic cells (granule cells, unipolar brush cells) expressing *SLC17A6,* VZ progenitors (*PTF1A*) differentiating into GABAergic cells (Purkinje cells, GABAergic interneurons) expressing *GAD1* and *GAD2*, as well as the noradrenergic lineage expressing *LMX1B* and *PHOX2B* (Figures 5B, S5B-D).

**Figure 5.**
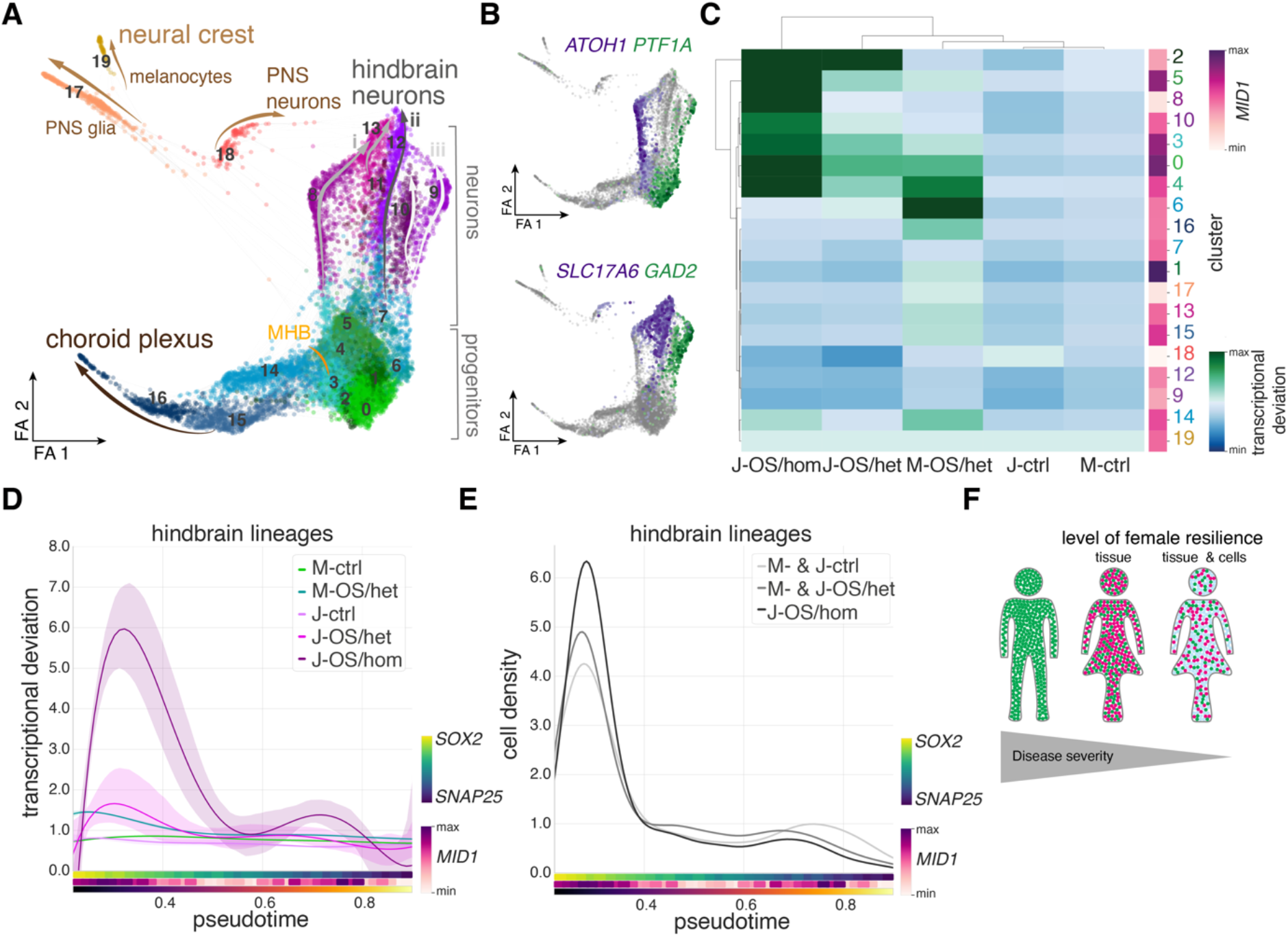
Molecular framework of OS phenotype. **A,** Force-directed graph embedding and Leiden clusters (indicated with numbers) of the neural cells (excluding clusters 1, 3, 11, 14, 16, 20 from the entire data set as shown in Figure 3A) highlighting various differentiation trajectories. Note the main three distinct neuronal hindbrain lineages i) glutamatergic neurons, ii) noradrenergic neurons, and iii) GABAergic neurons. Number of cells total: 11923, M-ctrl: 2017; M-OS/het: 4366; J-ctrl: 1565; J-OS/het: 615; J-OS/hom: 3360. **B,** Lineage reconstruction showing the mutually exclusive trajectories of rhombic lip progenitors (*ATOH1*) to glutamatergic cells (*SLC17A6*) and ventricular zone progenitors (*PTF1A*) to GABAergic cells (*GAD2*). **A, B,** FA refers to force atlas **C**, Heatmap showing the transcriptional deviation in each cluster from the respective control line and the corresponding *MID1* expression level. The dendrogram clustering the samples separates them according to their genotype with heterozygous lines showing intermediate deviation. There is no apparent correlation between the level of *MID1* expression and the degree of transcriptional deviation. **D**, Plot showing the pseudotemporal ordering of the transcriptional deviation in each cell along the hindbrain differentiation trajectory in different genotypes with the 95% confidence interval as a transparent band of the same color in the background. For orientation purposes the plot includes *MID1* expression as well as the *SOX2*/*SNAP25* ratio (NPCs versus neurons) averaged in 50 equally sized pseudotemporal bins. Besides the intermediate transcriptional deviation in heterozygous lines this plot reveals that the divergence in both heterozygous and homozygous lines peaks at NPC stages. **E,** Density distribution plot highlighting the accumulation of mutant transcriptomes at NPC stages. **F,** Schematic highlighting that resilience of female tissues is enhanced through the possibility to reactivate genes on the second X- chromosome, thereby ameliorating the phenotype of X-linked NDDs.

To elucidate the molecular framework driving the early disruption of the balance between proliferation and differentiation observed in OS brain organoids (Figure 4), differentially expressed genes in NPCs (clusters 0, 1, 2, 3, 4, 5, Figure 5A) between female ctrl and OS lines were determined followed by GO analysis (Table S2, Figures S5E-G). Amongst the differentially expressed genes, we found *ZIC1* and *ZIC2* (Figure S5E). Mutations in *ZIC1* and *ZIC2* lead to Dandy-Walker syndrome (Grinberg et al., 2004) characterized by malformation of the cerebellum. Interestingly, similar to *MID1* (Figure 4) these genes regulate neuronal differentiation (Aruga et al., 2002). Hence, our data suggest a partial overlap and a surprising molecular crosstalk linking OS and Dandy-Walker pathogenesis. Moreover, compared to wildtype, OS NPCs showed *MID1* genotype dependent (Figure S5H) gradual deregulation of more general cell cycle regulators associated with proliferation and differentiation (e.g. *CDK6, CCND1, KIF2A, NIN, CDC42,* Figures S5E-H) (Broix et al., 2018; Cappello et al., 2006; Grison et al., 2018; Lange et al., 2009; Shinohara et al., 2013), suggesting that altered cell cycle progression accounts for the accumulation of NPCs at the expense of neurons in OS brain organoids (Figure 4).

To directly assess whether altered cell cycle dynamics contribute to the development of the OS phenotype, BrdU pulse-chase experiments were conducted in d30 brain organoids (Figure S5I). We determined the number of postmitotic cells (Ki67^-^) that had incorporated the thymidine analog BrdU (BrdU^+^, i.e., passed through S-phase) during the time of the pulse-chase experiment. Quantification of BrdU^+^ Ki67^-^ cells within the organoid VZLS showed a reduced cell cycle exit of NPCs in OS organoids and a concomitant decrease of cells that differentiated into neurons (BrdU^+^ NeuN^+^) (Figures S5J-N). In addition, more BrdU^+^ cells were retained within the VZLS (Figure S5O) correlating with the observed increase of the VZLS area in OS organoids (Figure 4A). These data indicate that MID1 controls the balance between proliferation and differentiation of NPCs with hindbrain identity.

### Molecular phenotype attenuation on a cellular level

The severity of the phenotype in OS brain organoids (Figure 4) varied depending on the presence (M-OS/het, J-OS/het) or absence (J-OS/hom) of a wildtype *MID1* allele on the inactive X-chromosome with heterozygous lines showing intermediate phenotypes. To disentangle these tissue level phenotypes with cellular resolution we further leveraged the scRNA-seq data and computed the transcriptional deviation of each cell from their respective controls. These analyses revealed that in particular cells in the cerebellar clusters (0, 2, 3, 4, 5, 6, 8, 10) differed strongest while the transcriptomes of midbrain (14), choroid plexus (15, 16) and neural crest (17, 18, 19) derived cells varied less upon *MID1* mutation (Figure 5C) with slight variations between M- and J-lines. Importantly, lines with a wildtype *MID1* allele on the inactive X-chromosome (M-OS/het, J-OS/het) exhibited an intermediate transcriptional phenotype reminiscent of the characterization of tissue composition (Figures 4A-D, S4E, G). To temporally resolve the impact of *MID1* mutation on the NPC-to-neuron differentiation trajectory of cerebellar lineages we plotted the degree of transcriptional deviation along the developmental pseudotime (Figure 5D). The deviation from controls of both, heterozygous and homozygous lines was strongest early in the lineage at the NPC stage with high expression of the NPC marker *SOX2* and low expression of the mature neuronal marker *SNAP25*. These transcriptional differences occurred concomitant with an accumulation of cells at a transcriptional NPC stage in OS lines (Figure 5E) in accordance with the increase of VZLS and SOX2 in OS organoids (Figure 4). The observation that cells with homozygous *MID1* mutations (J-OS/hom) react stronger to the lack of functional *MID1* and heterozygous lines (M-OS/het, J-OS/het) show intermediate transcriptional phenotypes uncovers the gradual manifestation of the phenotype on an individual cell level. Our data show that the reactivation of the expression of alleles from the inactive X-chromosome leads to a dynamic increase in the diversity of the available allele pool. Furthermore, our data demonstrate that in NDDs caused by X-chromosomal mutations, this mechanism can enhance female brain resilience beyond the general random X-inactivation in tissues to ameliorate the phenotype on an individual cell level (Figure 5F).

## Discussion

Escape from X-inactivation resulting in biallelic expression and often to expression bias in female compared to male cells is a prime candidate molecular mechanism for sexual dimorphism in disease development. Genes escaping X-chromosomal inactivation have been identified employing interspecies hybrids (Cotton et al., 2013), in sequencing approaches using SNPs (Andergassen et al., 2017; Carrel and Willard, 2005; Tukiainen *et al*., 2017; Wainer Katsir and Linial, 2019), by analyzing DNA methylation (Cotton et al., 2015; Schultz et al., 2015) or by combining several approaches *in silico* (Sauteraud et al., 2021). In humans however, these studies were impeded by the difficult access to primary material and often restricted to readily accessible cells such as fibroblasts and lymphocytes or postmortem material resulting in analytical snapshots representing single time points disregarding developmental trajectories. In an experimental set-up combining cell reprogramming and single iPSC clone selection with *in vitro* neural differentiation protocols into NPCs and neurons (2D) or brain organoids (3D) and allele-specific analyses of bulk- and scRNA-seq data, we were able to follow mono- and biallelic expression of X-chromosomal genes across differentiation. We identified three different classes of escape genes: (i) Full escapees that escaped inactivation already in iPSCs and kept their escapee status throughout all differentiation stages. These largely overlapped with genes that had been described as constitutive escapees before and are thought to be able to avoid becoming part of the heterochromatin of the inactive X-chromosome (Carrel and Willard, 2005; Fang *et al*., 2019; Tukiainen *et al*., 2017). (ii) Reactivating genes exhibiting monoallelic expression in iPSCs that later during neural development induce expression from the inactive X-chromosome in a highly specific manner in terms of developmental time and lineage. (iii) Late-silencing genes, genes that were found to escape from XCI in the iPSCs, but during neural development silence one allele resulting in monoallelic expression at later timepoints. Our data strongly suggests that constitutive and facultative escapism are distinct entities and provide evidence that reactivation and late-silencing are possible mechanisms leading to facultative escape in a tissue-, lineage- and developmental time-specific manner.

The spatial organization of full escape, reactivating and late-silencing genes along the X-chromosome differs. Constitutive escapees were found to cluster particularly on the short arm of the X-chromosome (Carrel and Willard, 2005; Lahn and Page, 1999; Ross *et al*., 2005; Tsuchiya et al., 2004; Tsuchiya and Willard, 2000). This is related to X-chromosome evolution and the assumptions that the short arm of the X-chromosome has developed from an autosomal region distinct from the long arm (Ross *et al*., 2005) and that it is influenced by the pseudoautosomal region 1 (PAR1), that is homologous to Y-chromosomal sequences and as a whole constitutively escapes X-inactivation (Balaton and Brown, 2016; Posynick and Brown, 2019). While we detected a higher number of full escapees on the short arm of the X-chromosome, the linear proximity between these genes was not different compared to a random background distribution. This might be due to a bias in the analysis that depends on expressed SNPs in the used cell systems, which makes some genes inaccessible for the analysis. However, the pair-wise distance between reactivating genes was lower than expected by chance suggesting clustering rather than random distribution of these genes. We found clusters of reactivating genes on both arms of the X-chromosome. In addition to the chromosomal localization, we assessed the epigenetic landscape of the reactivating genes compared to genes which had not been detected to show biallelic expression along the neural differentiation trajectory. Strikingly, we found reactivating genes to be associated with an epigenetic pattern of active transcription and genic enhancer states. This is in contrast to genes that were found silenced on the inactive X-chromosome throughout all differentiation states. Environment of these genes was found enriched for repressed chromatin states.

This data suggests that facultative escapism is achieved by a chromatin status allowing dynamic locus-, lineage- and developmental time-dependent reactivation and late-silencing of X-chromosomal genes induced by differentiation. It further draws the surprising picture that switching between mono- and biallelic expression of X-linked genes is a frequently used, fine-tuned mechanism of gene expression regulation during female neural development. Interestingly, within the group of reactivating genes but not full escapees and late-silencing genes we found a surprising overrepresentation of NDD associated genes beyond the known overrepresentation on the X-chromosome linking reactivating genes to the development of NDD phenotypes.

NDDs present with a substantial gender bias with females being less frequently and less severely affected than males (Pearse and Young-Pearse, 2019). X-inactivation and escape have been suggested to play a role in phenotype development of NDDs (Brand *et al*., 2021). Our findings suggest that particularly reactivating genes were equipped with the ability to dynamically switch between mono- and biallelic expression and further that reactivation of gene expression from the inactive X-chromosome influences NDD phenotypes. To test the functional implication of the reactivation of X-linked genes on NDD phenotypes, we have here used brain organoids to model the disease etiology of OS, an X-linked NDD caused by mutations in the *MID1* gene. We have compared brain organoids derived from female iPSC clones that exclusively expressed mutant *MID1* (the wildtype allele was silenced on the inactive X-chromosome) with male cells carrying the same loss-of-function mutation in *MID1*. We found that the derived female neural tissue showed milder phenotypes than mutant male tissue suggesting a protective mechanism in female cells. We could further show that indeed reactivation of the wildtype *MID1* allele from the inactive X-chromosome during neural differentiation underlies the attenuation of the phenotype in female, heterozygous tissue. This data suggests that reactivation of the second allele from the inactive X-chromosome is a potent mechanism for female cells to curtail NDD phenotype development and increase brain resilience, which will have to be considered in the future for the prognosis of X-linked NDDs in females. Also, the reactivation of mutant alleles from the inactive X-chromosome might have the potential to influence disease development. Severe X-linked *HUWE1*- associated encephalopathy for example has been found to develop despite preferential inactivation of the mutated X-chromosome (Moortgat et al., 2018). Reactivation of alleles carrying gain-of-function *HUWE1* mutations from the inactive X-chromosome in the brain could be the riddle’s solution here.

Our data furthermore implies that X-linked gene reactivation not only influences X-chromosomal NDD phenotypes, but through their gene- and protein interaction partners the reactivating genes impact on a protein network with more global implications for the cells. In a GO enrichment analysis of the networks interacting with the reactivating genes we found an overrepresentation of terms involved in two central control mechanisms: gene expression control on an epigenetic level on one hand and protein function regulation through the ubiquitin system on the other hand. Both systems are sensitive to dose changes (Barakat et al., 2010; Froyen et al., 2012; Jonkers et al., 2009; Smith et al., 2011; Zhang et al., 2020). Changes of expression dose through reactivation of the silent allele on the inactive X-chromosome will thus interfere with the interaction network. It is therefore conceivable that through a complex symphony of interactions conducted by the X-reactivation network the epigenetic machinery in female cells allows to modulate the expression not only from gonosomes but also autosomes. NDDs are frequently caused by single dominant heterozygous mutations or by the sum of several common variants with smaller disease effects in autosomal genes (Bourgeron, 2016). In mouse models, we and others have shown that through exogenous epigenetic modulation expression from the intact allele in diseases caused by heterozygous autosomal mutations can rescue disease phenotypes (Cooper et al., 2020; Kim et al., 2017). This raises the concept that the X-reactivation network installs a safety net in the developing female brain increasing resilience towards NDDs and could be part of a mechanistic explanation for higher mutational burden in females required for the clinical manifestation of NDDs (Jacquemont *et al*., 2014). For the first time we here describe X-chromosomal reactivation induced by neural differentiation. It will be interesting to see if other than differentiation signals can stimulate X-chromosomal reactivation. Depending on quality and quantity of such stimuli it is conceivable that reactivation may vary individually. It is attractive to hypothesize that individually differential reactivation has graded influence on NDD phenotype development and might therefore be a fundamental reason underlying clinical variability of NDDs.

Taken together, our data describe a hitherto undiscovered mechanism of differentiation-dependent gene regulation in female brain cells leading to a dynamic increase in the diversity of the available allele pool and having the potential to modulate NDD phenotypes.

## Acknowledgements

We thank Dandan Han for help with library preparation, the Helmholtz NGS core facility for sequencing, Magdalena Götz (BMC LMU Munich) and Michael Wegner (FAU Erlangen-Nürnberg) for sharing equipment and lab space, and Claudia Keller-Valsecchi (IMB Mainz) for valuable comments on the manuscript. This work was supported by grants from the German Research Foundation (BE 4182/7- 1; CRC1080, project number 221828878), core funding to the Francis Crick Institute from Cancer Research UK, The Medical Research Council and the Wellcome Trust (FC001002), Wellcome Trust (206410/Z/17/Z), and the ReALity excellence initiative of the University Mainz to B.B., by the CRC 1080 and the German Research Foundation (SCHW 829/7-1 and BE 41882/7-1) to Su.S. and BB, German Research foundation (KA3125/2-1; GFK2162/2 TP C2), Schram foundation (T287/29577/2017) to M.K., the Bavarian State Ministry of Sciences, Research, and the Arts (ForInter; F.2-F2412.30/1/24) to M.K. and S.F., Interdisciplinary Center for Clinical Research (IZKF) at the University Hospital of the FAU Erlangen-Nürnberg to M.K. (P068; Jochen-Kalden funding programme N7) and S.F. (P074).

## Author contributions

B.B., S.F., Su.S., M.K. conceived the study and designed the experiments. S.K., M.B., E.W., J.K. generated and engineered iPSC lines. P.L. provided patient fibroblasts of the M-lines. B.W. and J.W provided initial training for iPSC generation and cell lines. S.K., M.B., performed the 2D differentiation assays, QUASEP, western blot and (q)RT-PCR and analyzed the data. Je. W. co-supervised S.K.. M.B. processed the bulk RNAseq samples. D.H., and S.D., preprocessed the data and performed variant calling while H.T. performed the bioinformatic analyses under the supervision of S.G. and with input from K.L.. R.M., E.G., Sa.F., H.B., B.L. performed the organoid experiments and analyzed the data together with S.F. and M.K. The single cell RNAseq data was processed and analyzed by S.F. and M.K.. B.B., S.F., Su.S., and M.K. wrote the manuscript, with all authors contributing corrections and comments.

## Data availability

The bulk RNAseq data is deposited in the Sequence Read Archive (SRA) under the BioProject PRJNA819272. The scRNA-seq data used in this study will be deposited in the Gene Expression Omnibus (GEO). The data that support the findings of this study are available from the corresponding authors upon reasonable request.

## Code availability

The computational code used within this study is available upon reasonable request.

## Additional information

**Correspondence and requests for materials** should be addressed to S.F., Su.S. or M.K..

## Supplemental information

### Supplementary Figures and Figure legends

**Figure S1 Related to Figure 1 and Figure 2.**
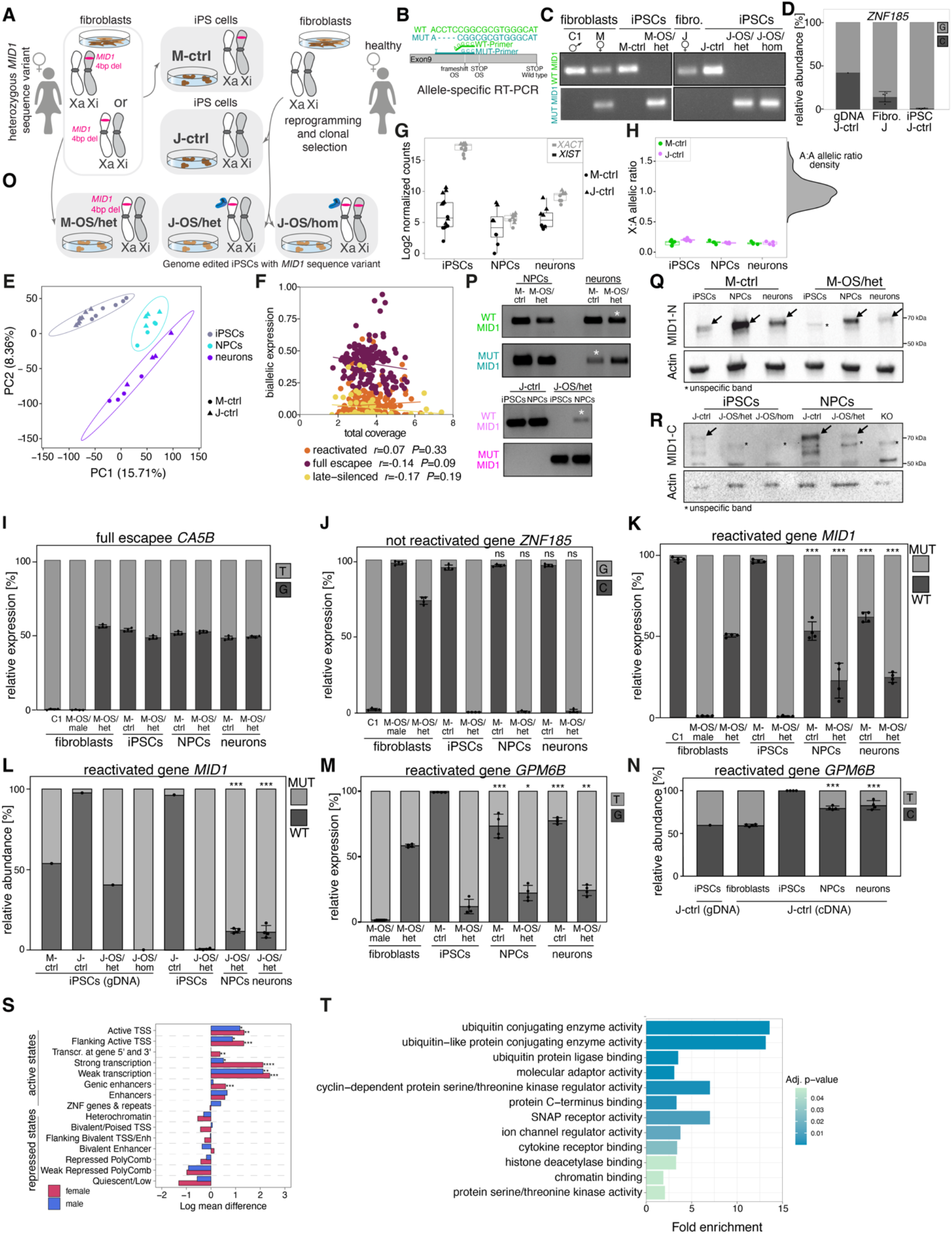
Characterization of X-reactivation. **A**, Scheme depicting the logic of iPSC clone selection, carrying heterozygous sequence variants in X-chromosomal genes, producing distinct iPSC lines from the same female donor with either one or the other X-chromosome active. **B**, Primer sequences used for allele-specific RT-PCR. **C,** Allele-specific RT-PCR analysis of *MID1*-activation status in fibroblasts and M iPSC lines (M-ctrl, M-OS/het) and J iPSC lines (J-ctrl, J-OS/het, J-OS/hom) derived from female fibroblasts (M, J). The C1 male control iPSC line shows the specificity of the mutant RT-PCR. **D,** Relative allele-specific expression of *ZNF185* transcripts in J-line cells analyzed by QUASEP. **E,** Principal component analysis of allelic ratios including 95% confidence ellipses confirmed a unique allele-specific expression profile for iPSCs, NPCs and neurons in the M-ctrl and J-ctrl cell lines. **F**, Scatterplot showing biallelic expression versus total coverage at the variant site for all reactivated, full escapee, and late-silenced genes. Regression analysis reveals no correlation between expression level and biallelic expression. **G,** Boxplots with log2-transformed normalized expression of *XIST* and *XACT* revealed a stable expression of both genes in all cell types of the M-ctrl and J-ctrl lines. **H**, Boxplots showing the normalized number of biallelically expressed variant sites from the X-chromosome divided by the normalized number of biallelic variants from autosomes (X:A allelic ratio). The values for all cell types were in the range corresponding to a normal XaXi status (Bar *et al*., 2019). Furthermore, all X:A values were lower than the A:A allelic ratios whose density is shown on the right y-axis of the plot. **G**, **H,** Boxplots show median, quartiles (box), and range (whiskers). **E**, **G**, **H,** M-ctrl iPSCs: *n=*6; J-ctrl iPSCs: *n=*8; M-ctrl NPCs: *n=*4; J-ctrl NPCs: *n=*4; M-ctrl neurons: *n=*4; J-ctrl neurons: *n=*4. **I,** Relative allele-specific expression of *CA5B* transcripts in M-ctrl and M-OS/het fibroblasts, iPSCs, NPCs, and neurons analyzed by QUASEP. **J**, Relative allele-specific expression of *ZNF185* transcripts in M-ctrl and M-OS/het fibroblasts, iPSCs, NPCs, and neurons analyzed by using QUASEP. Note that *ZNF185* does not show biallelic expression in iPSCs, NPCs or neurons (exact *P*-value from left to right: 0.81, >0.99, 0.80, 0.99). **K,** Relative expression of mutant and wildtype *MID1* transcript in M-ctrl and M-OS/het fibroblasts, iPSCs, NPCs, and neurons analyzed by QUASEP (*P*-values from left to right: <1.0*10^-4^, <1.0*10^-4^, <1.0*10^-4^, <1.0*10^-4^). **L,** Relative abundance of mutant and wildtype *MID1* gene and transcript in M-ctrl, J-ctrl, J-OS/het and J-OS/hom iPSCs, NPCs, and neurons analyzed by QUASEP (*P*- values from left to right: 2.0*10^-4^, 2.0*10^-4^). **M,** Relative allele-specific expression of *GPM6B* in M-ctrl and M- OS/het fibroblasts, iPSCs, NPCs, and neurons analyzed by QUASEP (*P*-values from left to right: <1.0*10^-4^, 1.6*10^-2^, <1.0*10^-4^, 4.2*10^-3^). **N,** QUASEP analysis of the relative abundance of *GPM6B* gene and transcript in J- ctrl fibroblasts, iPSCs, NPCs, and neurons (*P*-values from left to right: <1.0*10^-4^, <1.0*10^-4^). **I, J, K, L, M, N,** The unrelated wildtype male control line C1 and the hemizygous mutant M-OS/male fibroblasts are used as controls. Dots represent independent samples; all iPSC (gDNA), J-ctrl iPSC cDNA n=1; all other samples *n*=4; testing for statistical significance was performed for the values in NPCs and neurons compared to the values in the corresponding iPSCs. One-way ANOVA followed by Tukey’s multiple comparisons test; values are mean ± SD; *\*P*<0.05, *\*\*P*<0.01, *\*\*\*P*<0.001, ns *P*>0.05. **O,** Graphical scheme depicting the CRISPR/Cas9-mediated genome-editing of a wildtype female iPSC line (J- ctrl) in which the 4-bp deletion sequence variant was introduced hetero- or homozygously into the *MID1* gene (J- OS/het and J-OS/hom, respectively). M-OS/het iPSCs were selected for expression of the 4-bp deletion variant on the active X-chromosome. **P,** Allele-specific RT-PCR showing the expression of the mutant and wildtype allele in NPCs and neurons derived from M-ctrl and M-OS/het iPSC lines as well as in iPSCs and NPCs of J-ctrl and J- OS/het. **Q,** Protein levels of MID1 in M-ctrl and M-OS/het iPSCs, NPCs and neurons as determined by western blot using an N-terminal MID1 antibody. **R**, Western blot showing protein levels of MID1 in J-ctrl, J-OS/het, and J-OS/hom iPSCs and NPCs using a C-terminal MID1 antibody. An iPSC line with a CRISPR/Cas9-mediated removal of the complete *MID1* gene reveals that the band indicated by a star is unspecific. **Q, R** actin was used as a loading control. **S,** Chromatin state enrichment analysis between reactivated and inactive genes in female and male fetal brain samples. Positive log-mean differences correspond to higher levels of the corresponding state in reactivated genes and negative values indicate higher levels in inactive genes, exact *P*-values (top to bottom): 3.13*10^-2^, 7.78*10^-3^, 3.13*10^-2^, 1.42*10^-4^, 1.66*10^-3^, 3.13*10^-2^, 5.16*10^-5^, 3.73*10^-3^, 1.42*10^-4^, 3.51*10^-4^. **T,** Significantly over-represented GO terms in the PPI network of reactivated genes.

**Figure S2 Related to Figure 3.**
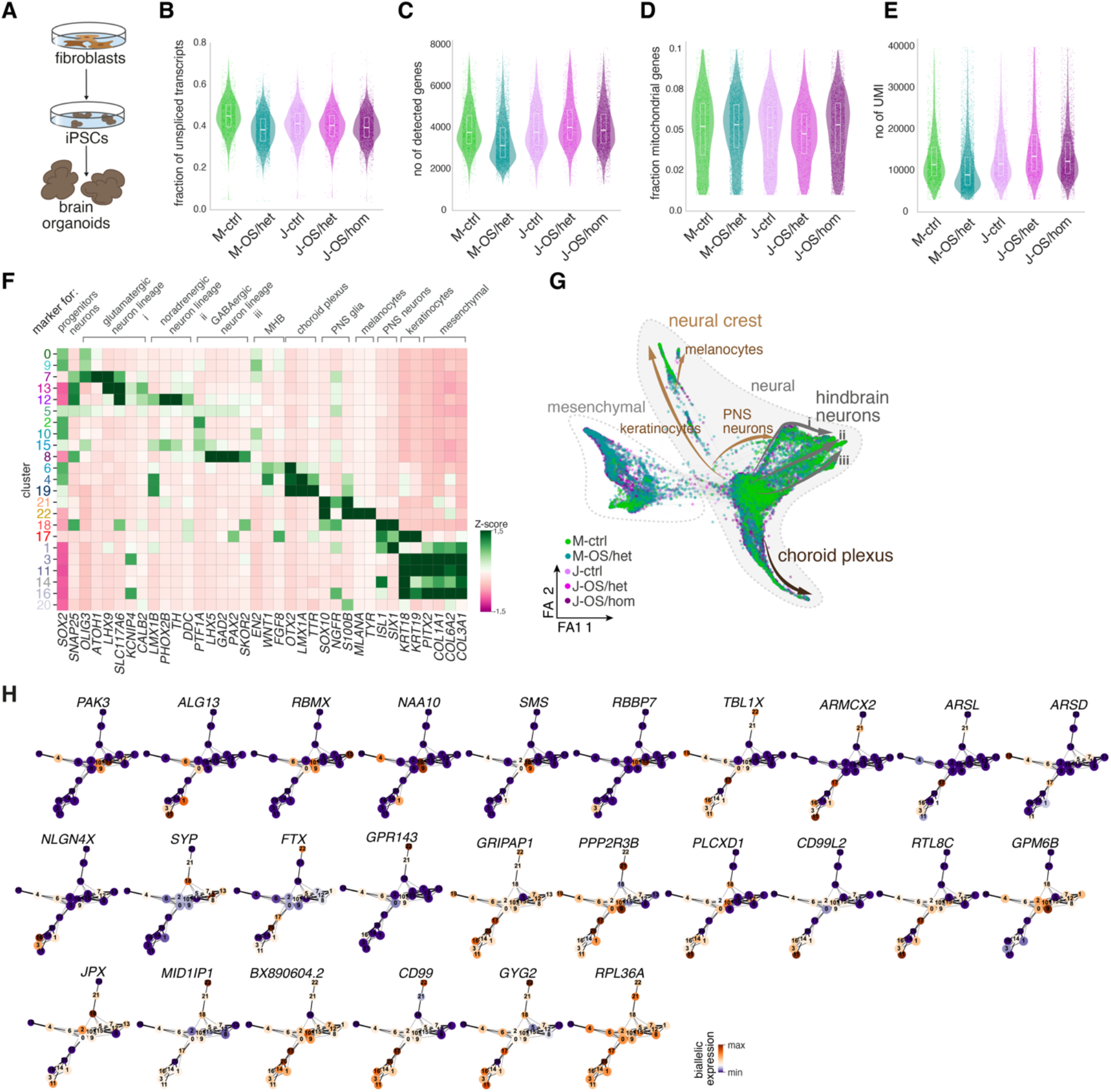
Allele-specific expression analysis of X-linked genes in brain organoids. **A,** Experimental scheme used to generate brain organoids. **B,** Violin plots depicting the fraction of unspliced transcripts in experimental conditions as determined by RNA velocity. **C, D, E,** confirm no difference in the quality of the scRNA-seq data of the different experimental conditions in terms of **C,** number of detected genes, **D,** fraction of mitochondrial genes, and **E,** number of detected UMIs. **F,** Heatmap showing marker gene expression discerning distinct cell types. **G,** Force directed graph embedding with experimental groups in different colors reveals similar contribution to all clusters and lineages. FA refers to force atlas **H,** Dynamic pattern of biallelic expression of detected genes on PAGA plots.

**Figure S3 Related to Figure 3.**
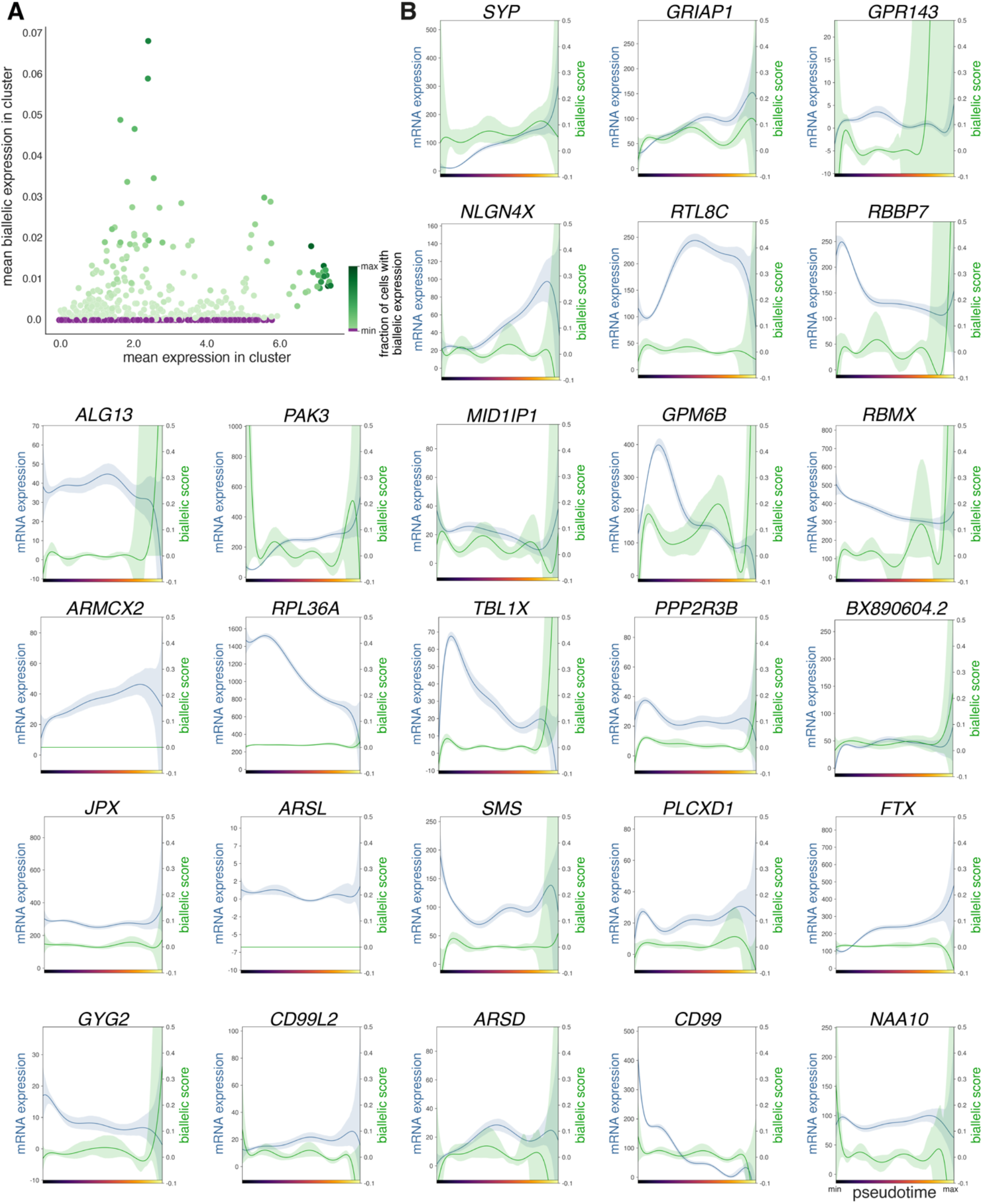
Biallelic expression and mRNA expression levels of biallelically expressed genes. **A,** Dotplot showing the mean total mRNA expression of the 26 detected reactivated genes in the 23 clusters (shown in Figure 3A) and the mean biallelic expression in the respective cluster. The color indicates the fraction of the cells exhibiting biallelic expression of the respective genes in the cluster. Note that there is no correlation between mRNA expression and usage of the second allele (Pearson correlation *r*=0.15). **B,** Total mRNA and biallelic expression of detected genes with dual allele usage along pseudotime of hindbrain development shows distinct behavior.

**Figure S4 Related to Figure 4.**
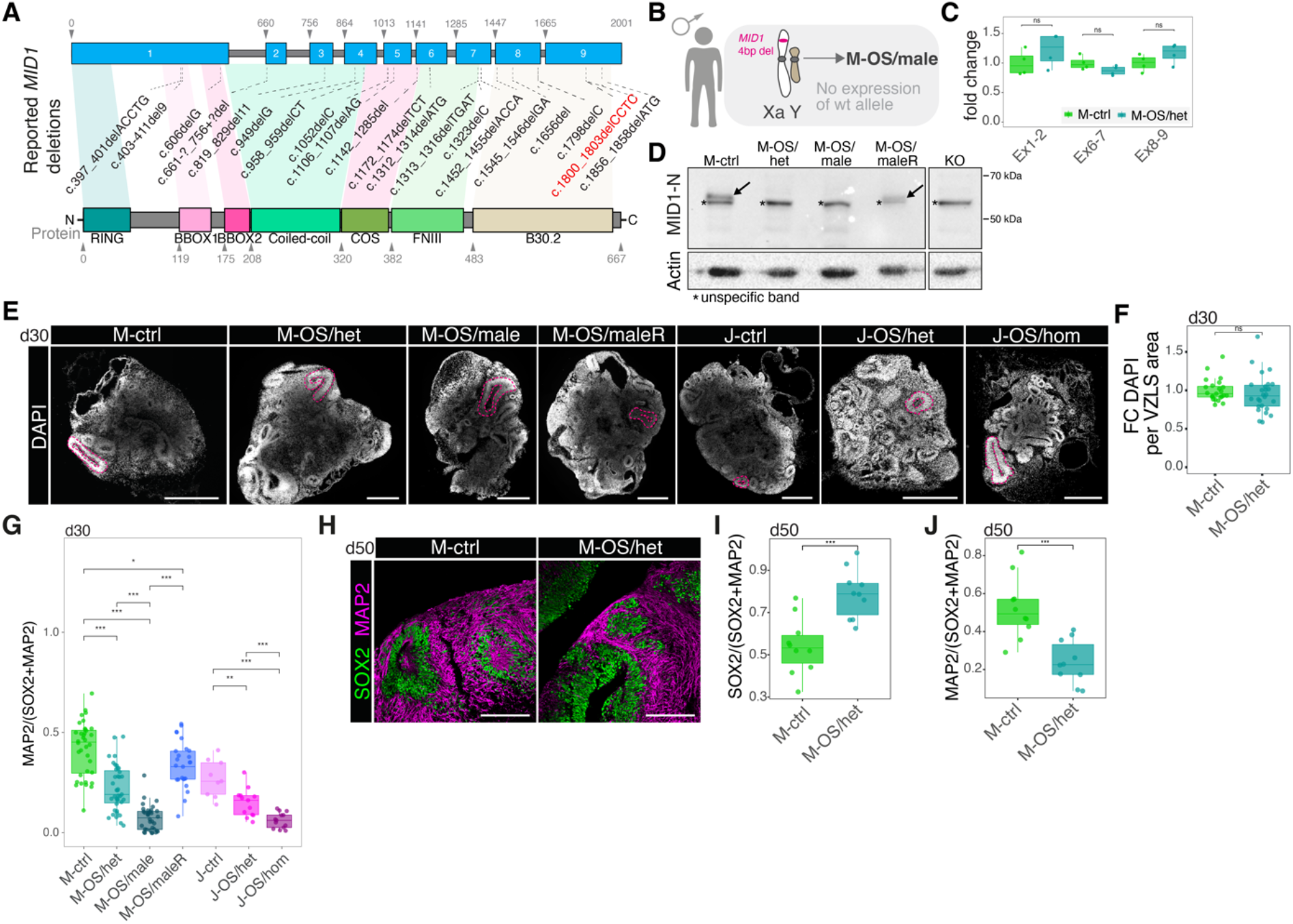
Characterization of brain organoids. **A,** Scheme indicating MID1 gene and protein and highlighting currently known mutations in the *MID1* gene including the 4-bp deletion c.1800_1803delCCTC studied throughout this work. **B,** Graphical depiction of the hemizygous 4-bp mutation in *MID1* in the M-OS/male iPSC line. **C,** qPCR was used to determine mRNA stability of the *MID1* transcript in different exons. Fold-changes were calculated over the mean of M-ctrl (M-ctrl, *n=*4; M-OS/het, *n=*4; exact *P* values (left to right): 0.2, 0.2, 0.34). **D,** Protein levels of MID1 in M iPSC lines as determined by western blot using an N-terminal MID1 antibody. An iPSC line with a CRISPR/Cas9-mediated removal of the complete *MID1* gene reveals that the band indicated by a star is unspecific. **E,** Micrographs depicting d30 organoid sections stained with DAPI. The pink dashed lines highlight examples of VZLS quantified in Figure 4A. Scale bar = 100 µm. **F,** Quantification of the number of DAPI nuclei per VZLS-area within brain organoid sections revealed no difference in cell density. Fold-change was calculated by dividing the cell density of each organoid by the mean cell density in controls (M-ctrl, *n=*23; M-OS/het, *n=*26; *P*=0.26). **G,** Quantification of the fraction of MAP2 in neural tissue (SOX2+MAP2+ area) on d30 (M-ctrl, *n=*36; M-OS/het, *n=*39; M-OS/male, *n=*34; M-OS/maleR, *n=*25; J-ctrl, *n=*9; J-OS/het, *n=*13; J-OS/hom, *n=*13; exact *P* values (top to bottom): 0.03, 9.7*10^-13^, 2.4*10^-9^, <2.2*10^-16^, 8.1*10^-9^, 7.9*10^-4^, 4.0*10^-6^, 4.3*10^-3^). **H,** Micrographs showing d50 brain organoid sections stained for SOX2 (green) and MAP2 (magenta). Scale bar=100 µm. **I,** Quantification of the fraction of SOX2 in neural tissue (SOX2+MAP2+ area) in d50 organoids (M-ctrl, *n=*10; M-OS/het, *n=*10; *P*=3.2*10^-4^). **J,** Quantification of the fraction of MAP2 in neural tissue (SOX2+MAP2+ area) in d50 organoids (M-ctrl, *n=*10; M-OS/het, *n=*10; *P*=3.2*10^-4^). **F**, **G, I, J,** dots represent individual organoids. **C**, **F, G, I, J,** boxplots show median, quartiles (box), and range (whiskers). Two-sided Wilcoxon rank sum test; *\*P*<0.05, *\*\*P*<0.01, *\*\*\*P*<0.001, ns *P*>0.05.

**Figure S5 Related to Figure 5.**
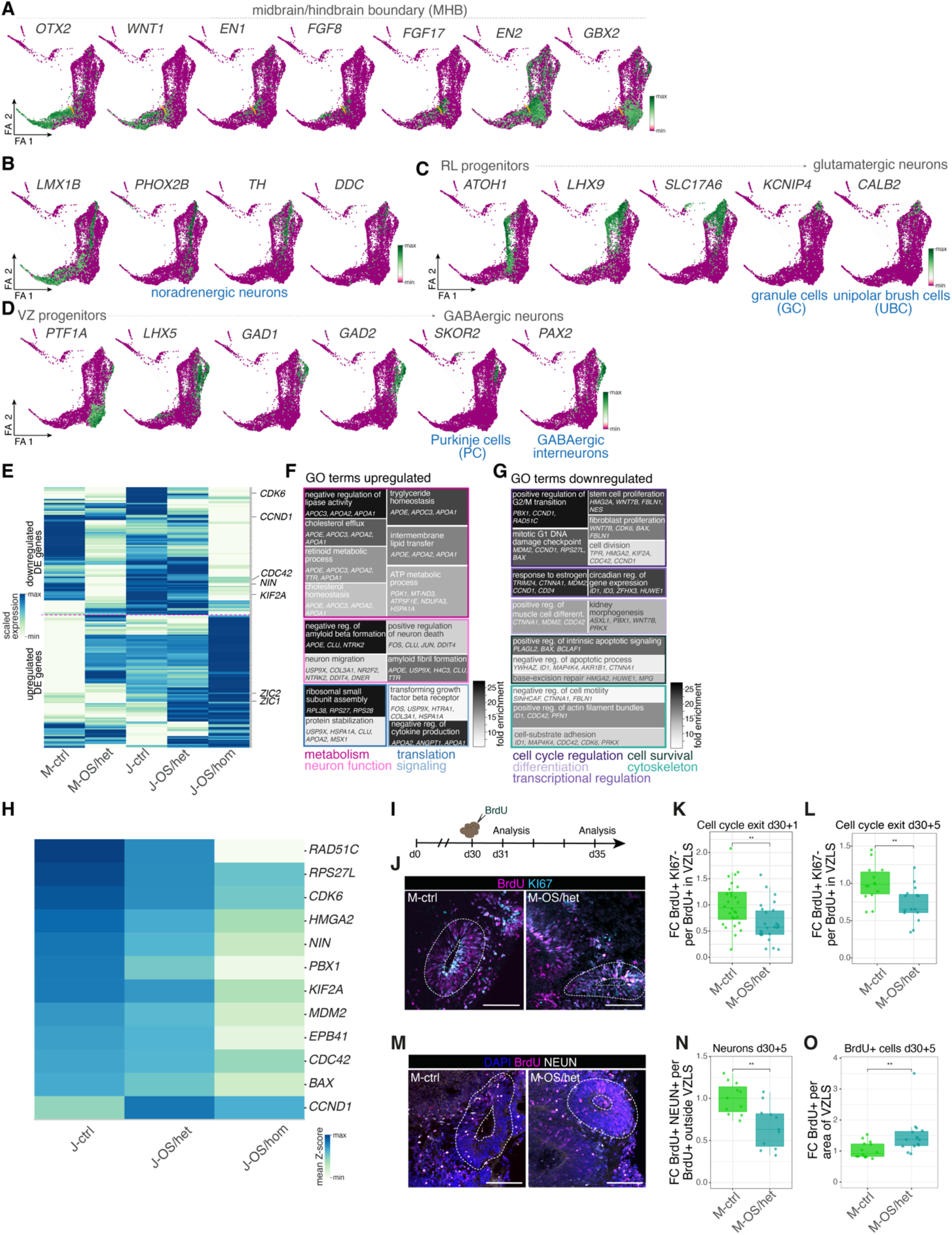
Molecular characterization of the OS phenotype. **A, B**, **C, D,** Force-directed graph embeddings colored by marker genes expressed **A**, at the midbrain/hindbrain boundary (MHB, yellow line), **B,** in the noradrenergic lineage, **C,** in the trajectory from rhombic lip (RL) progenitors towards glutamatergic neurons, and **D**, in hindbrain ventricular zone (VZ) progenitors giving rise to GABAergic neurons. FA refers to force atlas. **E,** Heatmap showing genes differentially expressed in NPCs (clusters 1, 2, 3, 4, 5, 6 in Figure 5A) between J-ctrl and J-OS/hom. Note an intermediate change in expression in J-OS/het compared to J-ctrl and J- OS/hom. **F, G**, Top 15 GO terms enriched in the differentially expressed genes in NPCs. GO terms were grouped as indicated below the tileplot. The color of the individual tiles indicates the fold enrichment. **H,** Heatmap showing expression of selected cell cycle related genes in ctrl and OS conditions. Note the intermediate expression pattern in J-OS/het versus J-OS/hom. **I,** Experimental scheme depicting the BrdU labelling experiment used to assess cell cycle exit of NPCs in d30 brain organoids. **J,** Micrographs showing representative stainings for BrdU (magenta) and KI67 (light blue) within VZLS (highlighted with white dashed line) 24 hours following BrdU pulse. Scale bar=100µm. **K,** Boxplot showing the quantification of the number of BrdU^+^ KI67^-^ cells per total number of BrdU^+^ cells quantified within VZLS 24 hours after BrdU addition. (M-ctrl, *n=*28; M-OS/het, *n=*26; *P*=1.7*10^-3^) **L,** Boxplot showing the quantification of the number of BrdU^+^ KI67^-^ cells per total number of BrdU^+^ cells quantified within VZLS 5 days after BrdU addition. (M-ctrl, *n=*14; M-OS/het, *n=*15; *P*=7.9*10^-3^) **M**, Micrographs of brain organoid sections stained for DAPI (blue), BrdU (magenta), and NEUN (white) 5 days following BrdU addition. VZLS are highlighted with white dashed lines. Scale bar=100 µm. **N**, Boxplot showing the quantification of BrdU^+^ NEUN^+^ cells per total BrdU^+^ cells quantified outside of VZLS 5 days following BrdU addition. (M-ctrl, *n=*11; M- OS/het, *n=*11; *P*=5.2*10^-3^). **O,** Boxplot depicting the quantification of BrdU^+^ cells per area of VZLS 5 days following BrdU addition. (M-ctrl, *n=*14; M-OS/het, *n=*15; *P*=3.2*10^-3^). **K, L, N, O** Fold-change was calculated by dividing the value for each organoid by the average value of controls in the respective batch; dots represent individual organoids; boxplots show median, quartiles (box), and range (whiskers); two-sided Wilcoxon rank sum test; *\*\*P*<0.01.

### Tables

**Table S1.** GO terms in the PPI network of reactivated genes.

**Table S2.** Differentially expressed genes in organoid-resident NPCs including up- and downregulated GO terms.

**Table S3.** Sequences of oligonucleotides used in this study.

## Methods

### Human fibroblast cell lines

Skin punch biopsies (4mm) were taken in the Children’s Hospital of the University Medical Center in Mainz. Primary human fibroblasts were isolated from skin punch biopsies as previously described with small adaptations (Vangipuram et al., 2013). Briefly, biopsies were cut in small pieces containing all skin layers and plated on a 6-well plate coated with 0.1% gelatin with fibroblast extraction media (FEM; DMEM (Thermo Fisher), 20% fetal bovine serum (FBS, Thermo Fisher), 1% penicillin/streptomycin (P/S) (Thermo Fisher)). Cells were monitored daily, and fresh medium was added every other day. Fibroblasts started to migrate out of the skin biopsies after 7-10 days and were transferred to two T75 flasks after 3-4 weeks using the standard splitting protocol with TrypLE™ Express enzyme (Thermo Fisher). Fibroblasts were then cultured in IMDM (Thermo Fisher), 15% FBS, 1% P/S. When reaching 90% confluency, fibroblasts were replated into T175 flasks and expanded as needed or frozen in liquid nitrogen for long-term storage. M-lines: Patients’ fibroblasts (for M-lines) had been established at Bristol Genetics Laboratory following the same procedure as described above.

### Reprogramming to pluripotent stem cells

Fibroblasts were reprogrammed into iPSCs with retroviruses by using the feeder-dependent approach of the CytoTune(TM)-iPS 2.0 Sendai Reprogramming Kit (Thermo Fisher) according to the manufacturer’s protocol. HEK 293T cells used for retrovirus production were cultured in IMDM, 10% FBS and 1% Pen/Strep and replated every three to four days at a ratio of 1:10 or 1:20 by using the standard splitting protocol using TrypLE ™ Express enzyme. To generate the retroviruses for reprogramming, HEK 293T cells were transfected by using Lipofectamine 3000 (Thermo Fisher) following the manufacturer’s instruction. Fibroblasts were transduced by retroviral spinfection. Briefly, 24 hours prior to transduction 1 x 10^5^ cells fibroblasts were seeded on a 6-well plate coated with 0.1% gelatin and cultured in IMDM plus 15% FBS. On day 0, medium was replaced with 5ml of retrovirus solution supplemented with a final concentration of 8μg/ml Polybrene (Sigma-Aldrich). Cells were then centrifuged for 1 hour at 800xg and the retrovirus solution was replaced by fresh medium afterwards. Treatment was repeated after 10 hours and after 24 hours. On day 2 (24 hours after the last treatment), cells were harvested and seeded in 10cm dishes on MEF feeder cells, one well of the 6-well plate per dish. On day 3, medium was changed to iPSC medium adding 0.1% SB431542 (StemCell Technologies). Medium was changed every other day. For the feeder-dependent approach of the CytoTune(TM)-iPS 2.0 Sendai Reprogramming kit, on day -2 fibroblasts were seeded in increasing concentrations on two 6-well plates (10-35 x 10^4^ cells/well) with reprogramming fibroblast medium (PFM; DMEM, 10% ESC-qualified FBS (Thermo Fisher), 1% NEAA (Thermo Fisher), 0.1% β-mercaptoethanol (Thermo Fisher)). On day 0, viral transduction was performed on 30-60% confluent cells using the Sendai virus-based reprogramming vectors containing the four Yamanaka factors, Oct, Sox2, Klf4, and c-Myc (Takahashi and Yamanaka, 2006). The required virus amount was calculated using the given formula from the manufacturer’s protocol and added to 1ml of RFM replacing the old medium. 24 hours after transduction, medium was replaced by fresh RFM. Cells were then fed every other day. On day 5 after transduction, MEF feeder cells were thawed and seeded on a 6-well plate (2.5 x 10^5^ cells/well) as described in the manufacturer’s protocol. On day 7 after transduction, fibroblasts were harvested, and seeded in different concentrations (5, 10, 20, 40, 80, 100 x 10^3^ cells/well) onto the previously prepared 6-well plate with MEF with RFM. On day 8, medium was replaced with iPSC medium (DMEM/F-12 (Thermo Fisher), 20% KOSR (Thermo Fisher), 1% NEAA, 0.1% β-mercaptoethanol, 1% P/S, 0.04% bFGF (Thermo Fisher)) and then changed every day. About 21-28 days after transduction, iPSC colonies were clearly emerging from the plate and approximately 50 colonies were picked and transferred to a single well of a 12-well plate coated with Matrigel (Corning) with mTeSR1 (StemCell Technologies). Medium was changed every other day; colonies were monitored regularly and expanded until ready for characterization. All iPSC lines were cultured in colonies in mTESR1 medium (StemCell Technologies) on Matrigel-coated dishes in 5% CO2 at 37°C until they reached a confluency of 80-90% and then replated at a ratio of 1:3 to 1:10 using a self-made enzyme-free splitting buffer containing PBS, NaCl and EDTA. Spontaneously differentiated cells were manually removed under a microscope. iPSC lines were maintained below passage 40 and karyotyped. All cells were regularly tested for the presence of mycoplasma using the PCR Mycoplasm Test Kit (PromoKine) or the LookOut Mycoplasma PCR detection kit (Sigma-Aldrich).

### Ethical approval

Patients’ fibroblasts (Liu et al., 2011; Schweiger *et al*., 1999; Trockenbacher et al., 2001) to generate M-ctrl, M-OS/het, M-OS/male, M-OS/maleR iPSCs were established in Bristol Genetics Laboratory, following consent for further analysis and usage for research in an anonymized way was given by the family. Fibroblasts to generate J-ctrl, J-OS/het, J-OS/hom, C1-male, and the KO-line iPSCs were taken at the University Medical Center in Mainz following approval by the local ethical committee (No. 4485). Consent for further analysis and usage for research in an anonymized way was given.

### Clonal selection of iPSC lines

Following reprogramming into iPSC, individual clones derived from the M- and J-fibroblasts, respectively, were picked and propagated for further analyses. Monoallelic expression of one X-chromosome in iPSCs was confirmed by using allele-specific PCR for the X-linked *MID1* gene (M-line) and by using a QUASEP assay for a heterozygous SNP located in the 3’untranslated region of the X-linked gene *ZNF185* (J-line).

### Genome editing

For CRISPR/Cas9 genome editing iPSCs were electroporated using the Lonza 4D-NucleofectorTM X Unit. 800.000 single cells per transfection reaction were pelleted. Cells were resuspended in 100μl electroporation buffer P3 (P3 Primary cell solution box, Lonza) plus 2.5μg of the CAG-Cas9-Venus plasmid (Yumlu et al., 2019) (pU6-(BbsI)sgRNA_CAG-Cas9-venus-bpA was a gift from Ralf Kuehn (Addgene plasmid # 86986) and 2.5µg of gRNA-containing plasmid. The CAG-Cas9-Venus plasmid did not contain any gRNAs, but the gRNAs were provided by using the gRNA cloning vector (Mali et al., 2013) (gRNA_Cloning Vector was a gift from George Church (Addgene plasmid # 41824)). Cell-plasmid solution was transferred to the electroporation cuvette (Amaxa^TM^ P3 primary cell 4D- Nucleofector^TM^ X, Lonza) and electroporated using the program CB-150 of the Nucleofector. After electroporation 100μl of RPMI (Thermo Fisher) was pipetted into the cuvette and incubated for 10 minutes at 37°C and then transferred to a fresh well of a Matrigel coated 6-well plate with 2ml of mTeSR1 supplemented with ROCK-inhibitor (10μM, StemCell Technologies). Single GFP-positive cells were sorted in a 96-well plate with mTeSR plus CloneR (StemCell Technologies). 14 days after plating, colonies were transferred to a 12-well plate with mTeSR plus CloneR. A small volume (∼10µl) of the dissociated cells was used for DNA isolation by using Quick-DNA Microprep Kit (Zymo research). The DNA was amplified by PCR and sequenced via Sanger Sequencing. Clones carrying the desired mutation were selected for further expansion.

### 2D iPSC differentiation into neuronal progenitor cells (NPCs)

iPSCs were differentiated into NPCs with the PSC Neural Induction Medium (NIM) (Gibco) (Havlicek et al., 2014). On day -1, iPSCs were washed with DMEM/F-12 and incubated with Collagenase IV (Thermo Fisher) for 20 min, followed by three wash steps with DMEM/F-12. Cells were scraped and resuspended with mTeSR1, transferred to ultra-low attachment plates and incubated for 24 hours. On day 0, medium was replaced with neuronal medium (NM: DMEM/F-12, 1% N2-supplement (Thermo Fisher), 2% B27-supplement (Thermo Fisher), 1% P/S) and replaced every other day. On day 7, EBs were seeded on one well of a 6-well plate coated with poly-Ornithine/Laminin (Thermo Scientific). Medium was changed every other day. On day 14, rosette-like structures were visible inside the attached EBs and were transferred to a new well of a poly-Ornithine/Laminin-coated 6-well plate. On day 2 after replating, medium was replaced and supplemented with 0.1% FGF2 (Thermo Fisher). NPCs were expanded as needed or frozen in liquid nitrogen for long-term storage. M-ctrl NPCs for bulk RNAseq were generated according to this protocol. All other iPSC-to-NPC differentiations followed a similar but commercial alternative: on day -1, iPSCs were seeded on a Matrigel-coated 6-well plate. After 24 hours, when cells reached confluency of 15-25%, medium was changed to Neural Induction Medium (NIM: Neurobasal (Thermo Fisher), 2% Neural Induction Supplement (Thermo Fisher), 1% P/S), renewed with increasing volumes every other day to compensate for cell growth. On day 7 of neural induction cells were replated following the manufacturer’s protocol in a Geltrex-coated (Thermo Scientific) 6-well plate (500x10^3^ cells/well) in Neural Expansion Medium (NEM: 49% Neurobasal, 49% Advanced DMEM (Thermo Scientific), 2% Neural Induction Supplement, 1% P/S) supplemented with ROCK-inhibitor (5μM, StemCell Technologies). After 24 hours, medium was exchanged with NEM and cells were monitored daily with medium changes every other day. When reaching confluency, cells were replated using TrypLE ™ Express and cultured on poly-Ornithine/Laminin-coated dishes with NM supplemented with FGF2 (20ng/ml).

### 2D Differentiation of NPCs into neurons

NPCs were seeded at low confluence (50-100 x10^3^ cells/well) onto poly-Ornithine/Laminin-coated cavities of a 6-well plate. 24 hours after seeding, cells were washed twice with PBS and medium was replaced by NM+VitA (DMEM/F-12, 1% N2-supplement, 2% B27+VitA supplement (Thermo Fisher), 1% P/S). Cells were cultured in a humidified incubator at 37°C and 8% CO_2_. Every 3-4 days, fresh medium was added to the cells. When reaching a total volume of 10-12ml per well, 20-50% of the medium was removed every time before adding fresh media. After 35 days, differentiated neurons were harvested as needed.

### Brain organoid formation

All iPSC lines were cultured in colonies in mTeSR Plus medium (StemCell Technologies) on Matrigel (Corning)-coated dishes in 5% CO_2_ at 37°C until they reached a confluency of 80-90%. Brain organoids were generated with slight modifications following the Lancaster protocol (Lancaster et al., 2013). Accutase (Gibco) was used to generate single cell suspensions of cells. Following centrifugation, cells were resuspended in organoid formation medium supplied with 4ng/ml of low bFGF (Peprotech) and 5µM ROCK-inhibitor (StemCell Technologies). Organoid formation medium consisted of DMEM/F12 + GlutaMAX-I (Gibco), 20% KOSR(Gibco), 3% FBS (Gibco), 0.1mM MEM-NEAA (Gibco), 0.1mM 2-mercaptoethanol (Sigma-Aldrich). 12,000 cells in 150µL organoid formation medium/well were reaggregated in low attachment 96-well plates (Corning) for at least 48 hours. After 72 hours half of the medium was replaced with 150µl of new organoid formation medium without bFGF and ROCK-inhibitor. At day 5 neural induction medium consisting of DMEM/F12 + GlutaMAX-I (Gibco), 1% N2 supplement (Gibco), 0.1mM MEM-NEAA (Gibco), and 1µg/ml Heparin (Sigma-Aldrich) was added to the EBs in the 96-well plate to promote their growth and neural differentiation. Neural induction medium was changed every two days until day 12/13, when aggregates were transferred to undiluted Matrigel (Corning) droplets. The embedded organoids were transferred to a petri dish containing organoid differentiation medium without vitamin A. Organoid differentiation medium consisted of a 1:1 mix of DMEM/F12 + GlutaMAX-I (Gibco) and Neurobasal medium (Gibco), 0,5% N2 supplement (Gibco), 0.1mM MEM-NEAA (Gibco), 100 U/ml penicillin and 100 µg/ml streptomycin (Gibco), 1% B27 +/- vitamin A supplement (Gibco), 0.025% insulin (Sigma-Aldrich), 0.035% 2-mercaptoethanol (Sigma-Aldrich). Three or four days later the medium was exchanged with organoid differentiation medium with vitamin A and the plates were transferred to an orbital shaker set to 30 rpm inside the incubator. Medium was changed twice per week. For fixation, organoids were transferred from petri dishes to 1.5ml tubes at day 30 and day 50. Organoids were washed with PBS and then fixed with 4% paraformaldehyde (PFA, Carl Roth) for 30 minutes. Time of PFA fixation was extended up to one hour depending on the size of the organoids. Afterwards, organoids were washed three times for 10 minutes with PBS and incubated in 30% sucrose (Sigma-Aldrich) in PBS for cryoprotection. For cryosectioning, organoids were embedded in Neg-50™ Frozen Section Medium (Thermo Fisher) on dry ice. Frozen organoids were cryosectioned in 30μm sections using the Thermo Fisher Cryostar NX70 cryostat. Sections were placed on SuperFrost Plus™ Object Slides (Thermo Fisher) and stored at −20°C until use.

### Immunohistochemistry

Cells on coverslips were fixed with 1ml of PFA (4% in PBS) for 20 minutes, washed three times with PBS, and incubated with blocking solution (PBS, 5% BSA, 0.3% Triton) for 1 hour. Afterwards, cells were incubated with primary antibodies diluted in blocking solution at 4°C overnight. On the second day, cells were washed three times with PBS, 0.1% Triton for 10 minutes, followed by an incubation with secondary antibody diluted in blocking buffer for 1 hour. After three wash steps with PBS, 0.3% Triton, coverslips were transferred to glass slides with 10μl of mounting medium (Vectashield, 0.5% DAPI, Vector Laboratories).

Organoid sections were post-fixed with 4% PFA for 15 minutes, washed three times for 5 minutes with 1 x PBS, and briefly washed with blocking solution containing 4% normal donkey serum (NDS, Sigma-Aldrich) and 0.25% Triton-X (Sigma-Aldrich) in 1 x PBS and subsequently incubated with blocking solution for at least one hour at RT. Tissue sections were incubated overnight at 4°C with primary antibodies diluted in antibody solution containing 4% NDS (Sigma-Aldrich) and 0.1% Triton-X (Sigma-Aldrich) in 1 x PBS. Following 3 wash steps using PBS with 0.5 % Triton-X (Sigma-Aldrich), secondary antibodies were diluted in antibody solution and incubated for one hour at RT. Finally, sections were washed three times for five minutes with PBS and PBS with 0.5 % Triton-X was used for the last wash step. Slides were counterstained with DAPI 1:1000 in PBS for 5 minutes, washed with PBS and mounted using Aqua PolyMount (Polysciences, Inc.). Slides were kept in a humidified chamber in the dark during the entire staining procedure.

Antibodies used were selected according to the antibody validation reported by the distributing companies. Mouse (IgG1) anti-MAP2 (Sigma-Aldrich; M4403; 1:300), mouse (IgG1) anti-PAX6 (Biolegend; 901301; 1:300), rat anti-BrdU (Abcam; ab6326; 1:300), rabbit anti-Ki67 (Invitrogen; MA- 14520; 1:300), rabbit anti-SOX2 (Abcam; ab137385; 1:300), mouse (IgG2b) anti-TUBB3 (Sigma; T8660; 1:300).The following secondary antibodies were used (1:500 dilution): goat anti-mouse IgG1 Alexa 488 (Thermo Fisher; cat.no. A21121), goat anti-mouse IgG Alexa 488 (Thermo Fisher; cat.no. A11001), goat anti-rabbit Alexa 488 (Thermo Fisher; cat.no. A11008), goat anti-rabbit Cy3 (Thermo Fisher; cat.no. A10520), goat anti-rat Alexa 555 (Thermo Fisher; cat.no. A10522), goat anti-mouse IgG1 Alexa 555 (Thermo Fisher; cat.no. A21127), goat anti-rat Alexa 633 (Thermo Fisher; cat.no. A21094), goat anti-rabbit Alexa 633 (Thermo Fisher; cat.no. A21070), goat anti-mouse IgG1 Alexa 633 (Thermo Fisher; cat.no. A21126).

### Microscopy and image analysis

Epifluorescence pictures were taken using AxioObserver (Zeiss), and the EVOS^TM^ M7000 Imaging System (Thermo Fisher). Confocal pictures were acquired at a LSM710 Confocal Microscope (Zeiss). Images were analyzed using FIJI (v1.52-1.53). To quantify the VZLS area we measured for each organoid section, the total organoid area (excluding areas covered by cysts), and the area covered by VZLS using FIJI (v1.52-1.53). In R (v3.5.1-4.1.2) we then calculated the fraction of organoid area covered by VZLS area, averaged these values from different sections from the same organoids and normalized the resulting values to the average value of the respective control organoids (J-ctrl for J-ctrl, J-OS/het and J-OS/hom; M-ctrl for M-ctrl and M-OS/het; M-OS/maleR for M-OS/maleR and M- OS/male) in each batch. To quantify the neural area covered by SOX2 and MAP2 in organoid sections we used OpenCV (v.4.4.0 - 4.5.1) in python (v3.9.1-3.9.10) for automated thresholding (same threshold for all pictures) and counting of thresholded pixels. The neural area was determined as the number of pixels thresholded for either SOX2 or MAP2 and the fraction of SOX2/neural area and the fraction of MAP2/neural area was calculated using NumPy (v.1.21.5) and pandas (v1.3.4).

### Western blot analysis

Protein lysates were generated from cell pellets using Magic Mix (48% urea, 15mM Tris pH7.5, 8.7% Glycerin, 1% SDS, 143mM β-mercaptoethanol) containing protease and phosphatase inhibitors (cOmplete Tablets easypack, PhosSTOP easypack, Roche) and transferred to a QIAshredder column (Qiagen). After centrifugation at 13500g for 2 minutes, the solution was transferred into a fresh tube and frozen at −80°C until needed. SDS gel-electrophoresis was used to separate proteins by their size. Proteins were transferred to a PVDF-membrane by using the Trans Blot Turbo Transfer Pack (Bio-Rad). Membranes were incubated for 24 hours with blocking buffer (PBS, 0.1% Tween, 5% milk), followed by 24 hours incubation with primary antibody diluted in blocking buffer. Membranes were washed 3 times for 10 minutes with PBS-T (PBS, 0.1 % Tween), incubated for 1 hour with secondary antibody diluted in blocking buffer, and washed 3 times for 10 minutes with PBS-T. Membranes were exposed using the Western Lightning Plus-ECL (Perkin Elmer) and imaging was performed by using ChemiDoc Imaging System (Bio-Rad). Images were prepared and analyzed using the Image Lab software (Bio-Rad). Mouse monoclonal anti-β-ACTIN (Sigma; A2066-200UL; 1:2000), rabbit polyclonal anti-MID1 C-terminal (Novus, NBP1-26612; 1:500), rabbit anti-MID1 N-terminal (courtesy of Dr. Sybille Krauss; 1:500) (Schweiger et al., 2017).

### qRT-PCR

Total RNA was extracted with the RNeasy Mini Kit (Qiagen) or the High Pure RNA Isolation Kit (Roche). Samples were stored at −80°C until use. The RNA concentration was measured using a Nanodrop (PeqLab) or Qubit4 (Thermo Fisher). cDNA was generated starting from 125ng up to 500ng of total RNA with the Maxima First Strand cDNA Synthesis Kit (Thermo Fisher) or the PrimeScript RT Master Mix (Takara). In each individual experiment, equal amounts of RNA were used for the generation of cDNA. The primers used are listed in Table S3. For qRT-PCR analysis, all samples were run in triplicates each with a reaction volume of 10μL, using the QuantiFast SYBR® Green PCR Kit (Thermo Fisher) or TB Green Premix Ex Taq II (Tli RnaseH Plus) Kit (Takara), 1μM primers and 1μL of cDNA. The reaction was performed in a QuantStudio 6 Flex Real-Time PCR System (Thermo Fisher) or the StepOne Plus Real-Time PCR System (Thermo Fisher) using the following amplification parameters: 5 minutes at 95°C, 40 cycles of 10 seconds at 95°C and 1 minute at 60°C. Data were analyzed using the ΔΔCT method as previously described (Livak and Schmittgen, 2001); expression levels were obtained normalizing each sample to the endogenous *GAPDH* control.

### Allele-specific RT-PCR

Specific primers were generated to bind to the wildtype or to the mutant (4-bp deletion) *MID1* allele (Table S3). For each sample, two reactions were performed, one with the reverse wildtype primer, and one with the reverse mutant primer, both using the same forward primer. After an initial denaturation step (95°C for 2 minutes), 35 cycles of denaturation step (95°C for 30 seconds), primer annealing step (70°C for 30 seconds), and elongation step (72°C for 30 seconds) were followed by a final elongation step (72°C for 5 minutes). The PCR products were analyzed by separation on a 1,5% agarose gel with EtBr.

### Quantification of allele-specific expression by pyrosequencing (QUASEP)

Amplification and sequencing primers (Table S3) were designed by using the PyroMark Assay Design Software (v2.0, Qiagen) for the region of interest, with one of the amplification primers being biotinylated. PCR reactions were performed for each sample with the amplification primers using 60°C as annealing temperature. Each sample was measured in triplicates. The PCR product was then used for pyrosequencing following the standard protocol of the PyroMark Gold Q96 Reagents kit (Qiagen). A sequencing cartridge was prepared with all four nucleotides, the enzyme, and the substrate mix for sequencing, with the volumes calculated by the Pyro Q CpG (Qiagen) software. The results were analyzed using the same software.

### Bulk RNA-sequencing

The library for RNA-seq experiments was prepared from 5ng of total RNA using the Ovation^®^ Solo RNA-Seq Library Preparation Kit (Tecan) following the manufacturer’s instructions. The library was denatured for sequencing by mixing 5μl of the 4nM library with 5μl of NaOH (0.2 M). After incubating for 5 minutes, 5μl of Tris buffer (200nM Tris-HCl, pH 7) were added. Finally, 985μl of prechilled HT1 were added. The sequencing run on a NextSeq 500/550 was performed as a paired-end run with 2 times 76 cycles and an expected output of 50 million reads per sample for allele-specific analysis.

### Bulk RNA-seq data pre-processing

Sequencing reads were demultiplex and base call (BCL) files were converted into Fastq files using bcl2fastq conversion software (v2.17.1.14, Illumina). Sequence adapters were trimmed and reads shorter than 6 bp were removed from further analyses using Cutadapt (v0.18) (Martin, 2011). Quality control checks were performed on the trimmed data with FastQC (v0.11.7). Read mapping of the trimmed data against the human reference genome and transcriptome (hg19) was conducted using the STAR aligner (v2.5.3) (Dobin et al., 2013). PCR duplicates were removed from the mapped reads using the Python script nudup.py (v2.3) provided from NuGen (https://tecangenomics.github.io/nudup/).

### Bulk RNA-seq differential expression analysis

Read counts of each gene were quantified using the Subread tool featureCounts (v1.6.2) (Liao et al., 2014). Differential gene expression analysis was conducted using the R package DESeq2 (v1.28.1) (Love et al., 2014) with an adjusted *P*-value < 0.1 as cutoff.

### Allele-specific expression analysis (ASE) of bulk RNA-seq data

For ASE, all bulk RNA-seq data from the M-ctrl and J-ctrl lines were processed with the commercially available NVIDIA Clara Parabricks Pipeline (v3.5) (https://www.nvidia.com/en-us/clara/genomics/). The build-in RNA pipeline rna_gatk was employed to align the fastq-files to the hg19 reference genome with the STAR (Dobin *et al*., 2013) aligner. After coordinate sorting and marking duplicates of the resulting BAM file, a base recalibration step was performed before variant calling with gatk Haplotype Caller (McKenna et al., 2010; Poplin et al., 2018). Analyses were restricted to the X-Chromosome by defining *-L chrX*. For joint genotyping, a genomic database was built with gatk GenomicsDBImport including all variants from all samples and subsequent genotyping of the X-chromosomal variants was performed with gatk GenotypeGVCFs. Gene names were annotated by the Ensembl Variant Effect Predictor (McLaren et al., 2016) before transforming the resulting VCF file to a table format with gatk VariantsToTable for smaller file size and further downstream analysis in R. We used R base functions to remove all variants that did not intersect known genes, multi- and monoallelic variants as well as intermediate and very long indels (>50 bp). Then we calculated the allelic ratio for each variant site by dividing the number of reads mapping to the reference allele (No. Ref) by the total number of reads covering the variant site (No. Total). For further processing, we set a threshold of at least 5 reads in all samples to consider a variant (SNP or indel) to be expressed. We then calculated average allelic ratios for each cell type (iPSCs, NPCs, and neurons) in the sequenced M-ctrl and J-ctrl lines, respectively, by summing up all reads mapping to the reference allele across all replicate samples from the corresponding cell type and cell line and dividing this number by the total number of reads covering the variant site. To detect reactivated genes which are characterized by monoallelic expression in iPSCs but biallelic expression in NPCs and/or neurons, we first identified all variant sites with an allelic ratio <0.025 or >0.975 in iPSCs. The allelic ratio was converted to an estimate of the probability of expression 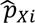 from the inactive chromosome Xi using the following formula

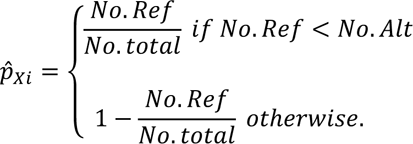

We then employed one-sided binomial tests to investigate 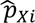 significantly greater than 0.025, indicating statistically significant expression from the inactive X-chromosome. *P*-values were adjusted for multiple comparisons using the Benjamini-Hochberg method implemented in the stats R package (v4.0.2). Results with an adjusted *P*-value<0.01 were considered statistically significant, indicative of a reactivated variant site. Variant sites were considered as escapee of X-chromosome inactivation 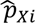 significantly greater than 0.1 in all cell types (iPSCs, NPCs, and neurons). To detect late-silenced ASE sites, we first identified all variants with 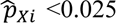 in NPCs and neurons and then used one-sided binomial tests to examine if Xi expression was significantly greater than 0.025 in iPSCs.

### Gene-wide estimates of biallelic expression

For genes that intersected only on ASE variant sites, we used the Xi expression for the respective variant as an overall estimate of biallelic expression for the whole gene. If a gene contained more than one type of biallelically expressed variants, the gene was assigned to a single category (reactivated, escapee or late-silenced) in a blinded fashion following a predefined set of rules: 1) the gene was assigned to the category with the highest number of biallelic variants 2) if a gene covered the same number of biallelic variants from different categories, then SNPs were considered as more reliable than indels, and variants of the same type with a higher coverage were preferred. After assigning genes to a unique category, we used the metaprop function from the meta R package (v4.18.1) to calculate gene-level estimates of biallelic expression from values for individual variants by using an inverse-variance method with logit-transformed proportions of Xi expression. 99% confidence intervals of biallelic expression based on a normal approximation were also obtained with the metaprop function.

### Manual curation of biallelically expressed variant sites

All predicted biallelic variant sites were subjected to manual inspection in a blinded fashion. 13 variants from the M-ctrl and 17 variants from the J-ctrl line were flagged as false positives because the predicted biallelic expression was due to all but one replicate expressing the reference allele and only one the alternative (or vice versa). These positions were then excluded from the analysis.

### Estimating the X:A allelic ratio

The X:A allelic ratio was calculated as reported before (Bar *et al*., 2019). First, we determined the number of biallelically expressed variant sites in each iPSC, NPC, and neuron sample having an Xi expression >0.1 with a Benjamini-Hochberg adjusted *P*-value <0.01 using a one-sided binomial test. We applied the same filtering criteria for autosomal variants as described for the X-chromosomal variants. To account for differences in the number of biallelic variants on each chromosome due to differences in gene density, we divided the sum of predicted biallelic variants by the number of genes on the respective chromosome based on the Ensembl annotation (https://genome.ucsc.edu/cgi-bin/hgTables). Subsequently, we calculated the X:A allelic ratio by dividing the normalized number of biallelic variants on the X-chromosome with the average of the normalized number of biallelic variants on the autosomes.

### X-chromosome ideograms

Ideograms showing the cytogenetic location of reactivated, escapee and late-silenced genes were produced with the karyoploteR package (v1.4.1) based on the hg19 genome assembly. Gene coordinates were obtained from the NCBI RefSeq annotation provided in the SMITE package (v1.16.0).

### Enrichment of NDD associated genes

We calculated the expected number of X-chromosomal genes associated with NDD by taking advantage of a recently published database review defining genes associated with NDD with high confidence (Leblond *et al*., 2021). We randomly sampled the whole genome, selecting the number of genes found on the X-chromosome and present in the database (*n*=836) for 10^6^ times and determine the number of NDD associated genes in each iteration. The mean value was determined as the expectation value and the fold change as the ratio of the NDD genes present on the X-chromosome / expectation value. The significance of observing a value higher than the measured value of NDD genes on the X-chromosome was determined by the cumulative distribution function. To determine the expectation value for reactivated, late-silenced and escapee genes on the X-chromosome being associated with NDD, we randomly sampled the 836 X-chromosomal genes selecting the number of genes present in each category (reactivated: *n*=56, late-silenced: *n*=16, escapee: *n*=38) for 10^6^ times and determined the number of NDD associated genes in each iteration. The mean value was determined as the expectation value and the fold change as the ratio between observed numbers of NDD genes / expectation values in each class. The significance of observing a value higher than the measured value of NDD genes in each class was determined by the cumulative distribution function.

### Pairwise distance analysis of biallelically expressed genes

We calculated the median pairwise distance for reactivated, escapee, and late-silenced genes, respectively. We tested the probability of observing a lower median pairwise distance by chance by comparing the actual distance to a background distribution of median pairwise distances for 1000 sets of randomly selected X-linked genes which were expressed in NPCs and neurons in our bulk RNA-seq data (>10 reads in each sample following DESeq2 normalization of the data). The *P*-value of observing a lower median pairwise distance by chance was obtained with the cumulative distribution function of the normal distribution.

### Protein-protein interaction network of reactivated genes

To construct a protein-protein interaction (PPI) network of reactivated genes, we employed the data provided by human binary reference interactome (HuRi) (Luck et al., 2020) at http://www.interactome-atlas.org/. The 42 protein-coding reactivated genes that were included in the HuRi portal, were used to create the network. To construct a differentiation-specific PPI, we filtered for genes that were expressed in all NPC and neuron samples in our bulk RNAseq data (>10 reads in each sample following DESeq2 normalization). We employed the igraph package (v1.2.6) in R to visualize the network. GO term analysis of the PPI network of reactivated genes was performed with the clusterProfiler package (v3.16.1) using an adjusted *P*-value cut-off of 0.05. Redundant GO terms were combined using the simplify function.

### Chromatin state analysis of reactivated genes

To investigate the association between X-chromosome reactivation and chromatin state, we employed the 15-state chromatin model from the Roadmap Epigenomics Consortium (Roadmap Epigenomics *et al*., 2015) using the female (sample E082) and male (sample E081) fetal brain samples. The annotation files for the epigenomes were downloaded from https://egg2.wustl.edu/roadmap/web_portal/chr_state_learning.html. We used the ChromDiff tool (Yen and Kellis, 2015) to estimate the percentage of each gene body corresponding to each of the 15 chromatin states in the two epigenome samples. We then compared the chromatin state profiles of the 53 reactivated genes that were included in the GENCODE annotation files distributed with the ChromDiff tool against the profiles of X-chromosomal genes expressed in NPCs and neurons (>10 normalized reads in all samples) but not showing biallelic expression in our assay and not being previously reported as escapee genes (Oliva *et al*., 2020). We then performed one-sided Wilcoxon rank sum tests to investigate the enrichment of epigenomic states in reactivated as well as genes not detected as biallelically expressed separately in the female and male fetal brain samples. *P*-values were adjusted for multiple comparisons with the Benjamini-Hochberg method and an epigenomic state was considered enriched in the respective gene category if the adjusted *P*-value was <0.05. To compare the enrichment of chromatin states in the female relative to the male fetal brain, we subtracted the male log-mean difference of enrichment of reactivated relative to inactive genes from the estimate in the female sample. We then transformed this difference to a fold change ratio using the following formula:

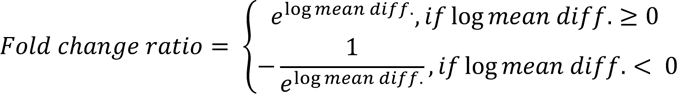

To evaluate the statistical significance of the fold change ratio, we produced a background distribution by randomly permutating the assignment of reactivated and inactive gene labels 1000 times and then calculating the fold change ratio in these samples for each epigenetic state. Statistical significance of the estimates was obtained by comparing the actual values to the background distributions based on the cumulative distribution function of the normal distribution. *P*-values were adjusted for multiple comparisons with the Benjamini-Hochberg method.

### Sex-biased expression of reactivated, escapee, and late-silenced genes

Information about sex biases in the expression of the biallelic and inactive genes predicted here was extracted from the study by Oliva and colleagues (Oliva *et al*., 2020) which investigated systematic differences of gene expression between males and females across 44 human tissues form GTEx V8. We compared the distributions of the number of tissues with a female bias in each of our gene categories using a Kruskal-Wallis test followed by Dunn’s test for pairwise multiple comparisons with Benjamini-Hochberg adjusted *P*-values as implemented in PMCMRplus v1.9.3 in R.

### Single cell RNA-seq data generation

For the 10X Genomics experiment, organoids were dissociated using the Neural Tissue Dissociation Kit P (Miltenyi Biotec). Briefly, selected organoids were cut into smaller pieces, washed with medium, and three times for 5 minutes with 1xPBS. Organoid pieces were transferred to a tube containing the enzyme mix P (according to the manufacturer’s protocol) and incubated at 37°C for 10 minutes. Pieces were then triturated gently with a 1000p pipette tip and incubated for another 10 minutes at 37°C in the presence of enzyme mix A (according to the manufacturer’s protocol). Pieces were then triturated gently with a 1000p and a 200p pipette tips and incubated for 5 minutes at 37°C. Cell suspension was filtered with a 30 µm filter (Miltenyi Biotec) and centrifuged at 300xg for 5 minutes. After a second filtration step with a 20µm filter (Miltenyi Biotec) and subsequent centrifugation as described above cell pellet was resuspended in 100µl 1xPBS (without Ca^2+^ and Mg^2+^). Cells were counted and tested for viability with Trypan Blue and the automated cell counter Countess (Thermo Fisher). Cells were diluted to an appropriate concentration to obtain approximately 5000 cells per lane on a 10X Next GEM chip G v3. Libraries were constructed according to the protocol of 10X genomics and sequenced on an Illumina NovaSeq 6000.

### Single cell RNA-seq data preprocessing, clustering, visualization

The functions count and aggr of the Cell Ranger software (10x Genomics. Version 4.0.0 – 6.0.2) were used to for de-multiplexing, aligning to the GRCh38 reference genome and for sequencing depth normalization. Scanpy (Wolf et al., 2018) (v.1.8.1) was used for further preprocessing, clustering, embedding, and visualization. Cells were excluded in which we detected less than 500 genes, less than 3000 UMI counts, more than 50000 counts, fraction of the transcriptome of less than 1% or more than 10% mitochondrial genes. Moreover, genes expressed in less than 5 cells were excluded from further analysis. Doublets were identified with Scrublet (Wolock et al., 2019) and filtered out. After normalization and log transformation, highly variable genes were determined using the default settings in Scanpy. Cell cycle scores, percentage of mitochondrial genes, and number of detected UMIs were regressed out to reduce their confounding effects. PCA analysis with the arpack wrapper in SciPy (v1.7.0) followed by determining the 15 closest neighbors in the top 20 PCs, batch integration using bbknn’s (v1.4.0) (Polanski et al., 2020) Euclidian metric and force-directed graph embedding utilizing ForceAtlas2 (Jacomy et al., 2014) provided by the python package fa2 (v.3.5) was used for embedding the transcriptomes. Clustering was performed with Scanpy’s python implementation of the Leiden (Traag et al., 2019) algorithm (v.0.8.7). For a coarse-grained mapping of clusters, their connectivity and lineages we applied the Scanpy implementation of the partition-based graph abstraction (PAGA) (Wolf *et al*., 2019).

### Single cell RNA-seq RNA velocity and pseudotime estimation

To determine spliced and unspliced transcripts, the Cell ranger produced BAM files were sorted by the cell barcode with samtools (v.1.10) and counted with velocyto (v.0.17) (La Manno *et al*., 2018). RNA velocity estimation was then done with scVelo (v.0.2.3) (Bergen *et al*., 2020). In detail, moments were calculated with the ‘connectivities’ mode on the top 50 PCs and 10 closest neighbors. After recovering the velocity dynamics, latent time was calculated, a measure for the developmental time (pseudotime) exclusively depending on transcriptional dynamics. RNA velocity was then computed, using latent time, differential kinetics and highly variable genes with the stochastic model.

### Differential gene expression and GO term analysis

Differentially expressed genes in NPC were determined between J-ctrl and J-OS/hom in the clusters 0, 1, 2, 3, 4, 5, and 6 of the neural lineages embedding using the rank_genes_groups function of Scanpy and testing for significance with the Wilcoxon rank sum test. Genes with an adjusted *P*-value < 0.01 and an absolute log2 fold change > 0.25 were considered differentially expressed. GO analysis was done with the R (v.4.1.2) package TopGO (2.44.0) considering genes expressed in NPCs as background and excluding GO terms with less than 5 annotated genes from the analysis. The default weight01 algorithm was applied to determine the GO enrichment. Only GO terms in which more than 3 differentially expressed genes were found and with a *P-*value < 0.05 as determined by Fisher’s exact test were considered.

### Transcriptional deviation

To determine the transcriptional deviation from control we determined for each cluster the differentially expressed genes between J-ctrl and J-OS/hom as well as M-ctrl and M-OS/het with *P*-value < 0.01 and an absolute log2 fold change > 2 (excluding cluster 11 due to the lack of cells from the J-OS/hom condition). For each differentially expressed genes we calculated the average expression in both controls (M-ctrl and J-ctrl) in this cluster. For each differentially expressed gene in each cell of the cluster we determined the fold change over the average expression of this gene in the corresponding control. For upregulated genes: cell^gene^/(average ctrl^gene^); for downregulated genes: 1/(cell^gene^/(average ctrl^gene^)). In each cell of the cluster, we determined the mean fold change of all cluster-specific differentially expressed genes as a measure for the transcriptional deviation. To determine the cluster-specific transcriptional deviation, we calculated the mean transcriptional deviation of all cells of that cluster in a cell line-specific manner. To resolve the temporal pattern of transcriptional deviation, the single cell values of transcriptional deviation were plotted against the RNA velocity based pseudotime estimation (latent time). The *MID1* expression levels as well as the ratio of *SOX2*/*SNAP25* expression were calculated by binning the pseudotime in 50 equally sized bins and calculating the mean values in J-ctrl cells.

### SNP calling and variant counting in scRNAseq data

To detect expressed heterozygous SNPs on the X-chromosome we used FreeBayes (v.1.3.5) (Garrison, 2012). The bam files for each line were used separately to detect intragenic variants with a PHRED quality score of 20. Variants with a sequencing depth normalized threshold of at least 50 reads per 5*10^9^ total reads were considered for further analysis (no. of total reads, M-ctrl:317987729; M-OS/het:273695358; J-ctrl:361543776, J-OS/het:485206581, J-OS/hom:396435551). The information on X-chromosomal variants from each line were merged and used for allele-specific alignments with the WASP (van de Geijn et al., 2015) implementation in the STAR aligner (v2.7.8a) (Dobin *et al*., 2013). Using NumPy (v.1.21.4) and pandas (v1.3.4) in python (v. 3.9.7), the number of UMIs for each variant in each cell was determined, and the absolute value of the natural logarithm of the ratio between the most abundant and the sum of all variants used as a measure for the allelic expression of a given gene in a given cell (monoallelic=0, biallelic with 50% transcripts from each allele=0.69). To calculate the degree of general biallelic expression from the X-chromosome in a cell, the values of biallelic expression for all biallelically expressed X-chromosomal genes were summed up.

To determine the usage of the second allele from the X-chromosome in each cell, we summed up the UMI counts of the less abundant variants (assuming the transcripts from the inactive X-chromosome are less abundant) and normalized the resulting sum with the total number of detected UMIs.

Cluster annotation to lineages were as follows according to the clusters in Extended Data Fig. 2e: hb-rl (hindbrain-rhombic lip): 0, 9, 7, 13; hb-vz (hindbrain ventricular zone): 2, 10, 8; nor (noradrenergic lineage): 12, 15; cp (choroid plexus): 4, 19; nc (neural crest): 18, 21, 22; mid (midbrain): 6; ker (keratinocytes): 17, meso: 1, 3, 11, 14, 16, 20.

### Statistics and reproducibility

Data were statistically analyzed with Microsoft Excel, GraphPad Prism or R using statistical tests indicated throughout the manuscript. No statistical methods were used to predetermine sample size. The investigators were not blinded to allocation and outcome analysis. The experiments were not randomized.

